# Dual Modes of Gene Regulation by CDK12

**DOI:** 10.1101/2025.09.22.677923

**Authors:** Yubao Wang, Apoorva Baluapuri, Cherubin Manokaran, Jing Ni, Tao Jiang, Roel C. Janssens, Jurgen A. Marteijn, Karen Adelman, Jean Zhao, Thomas M. Roberts

**Affiliations:** Departments of Cancer Biology, Dana-Farber Cancer Institute, Boston, MA 02115, USA; Department of Biological Chemistry and Molecular Pharmacology, Harvard Medical School, Boston, MA 02115, USA; The Eli and Edythe L. Broad Institute, Cambridge MA 02142, USA; Ludwig Center at Harvard, Boston, MA 02115, USA; Department of Molecular Genetics, Oncode Institute, Erasmus MC Cancer Institute, Erasmus University Medical Centre, Rotterdam, the Netherlands

## Abstract

The process of transcription is driven forward by the activity of kinases including CDK7, CDK9 and CDK12. Accordingly, acute inhibition of any of these kinases results in profound downregulation of gene expression. Here, we discover that loss or inhibition of CDK12 also significantly upregulates a set of coding and non-coding loci, whose activation could contribute to the anti-proliferative effects of CDK12 inhibitors. Mechanistically, CDK12 inhibition impairs transcription elongation, leading to increased RNA polymerase II termination or arrest in long genes. However, short genes such as MYC and enhancer RNAs are highly transcribed in the absence of CDK12 activity. Indeed, in HER2+ breast cancer, a malignancy where CDK12 is co-amplified with HER2 and its expression correlates with disease status, CDK12 inhibition markedly elevates MYC expression to induce lethality. The dual effects of CDK12 inhibition elucidated herein clarify its role in transcriptional control and have significant translational implications.

## Introduction

Gene transcription, a process by which RNA is synthesized by RNA polymerase II (Pol II) from DNA templates, is highly regulated, by a myriad of protein factors. In particular, the largest subunit of Pol II, RPB1, has an extended carboxy-terminal domain (CTD), consisting of an evolutionarily conserved heptad sequence (Tyr_1_-Ser_2_-Pro_3_-Thr_4_-Ser_5_-Pro_6_-Ser_7_) that is repeated 52 times in human cells. Distinct residues within the CTD are phosphorylated by the transcriptional kinases at specific stages of transcription, to regulate progression across the transcription cycle^1^.

Initiation of transcription is driven by transcription factors and co-factors, leading to recruitment Pol II at gene promoters. After synthesis of 30-50 nucleotides downstream of the transcription start site (TSS), Pol II machinery enters into a paused state, a stage critical for the control of gene expression^2^. CDK7, a component of the general transcription factor TFIIH, phosphorylates the CTD repeats at the Ser5 position during pausing, stimulating the association of RNA capping factors to protect the nascent RNA^2–4^. Pause release is catalyzed by the kinase P-TEFb (comprised of CDK9 and Cyclin T), and involves the phosphorylation of the Pol II CTD at Ser2 residues to enable the transition to productive elongation. CDK12 further phosphorylates Ser2 and the elongation machinery during transcription, to promote full length RNA synthesis^2, 5, 6^.

As CTD Ser2 phosphorylation is closely associated with transcription pause release and productive Pol II elongation, kinases that catalyze CTD Ser2 phosphorylation such as CDK9 and CDK12 have recently emerged as potential targets for cancer therapeutics^6–8^. From a mechanistic standpoint, acute inhibition of CTD Ser2 kinases – namely, inhibition that is brief enough to preempt any potential effects on cell cycle progression and cell survival – is expected to predominantly downregulate gene expression. Notably such inhibition particularly affects mRNAs with relatively short half-lives, a group of genes in which key oncogenic factors are overrepresented (Figure 1A). However, we have found that the alteration of gene expression observed upon CDK12 inhibition is drastically different from that resulting from CDK7 or CDK9 inhibition. Surprisingly, we observe rapid upregulation of a number of genes that could contribute to the efficacy of CDK12 inhibition in killing HER2+ breast cancer cells – an aggressive type of breast cancer in which the CDK12 gene is co-amplified with the HER2/ERBB2 gene. Detailed mechanistic assays suggest a model wherein CDK12 regulates an early elongation checkpoint. Loss of this checkpoint prematurely releases Pol II into elongation, which allows for upregulation of short genes and non-coding RNAs, while long messenger RNA genes are generally repressed.

**Figure 1.**
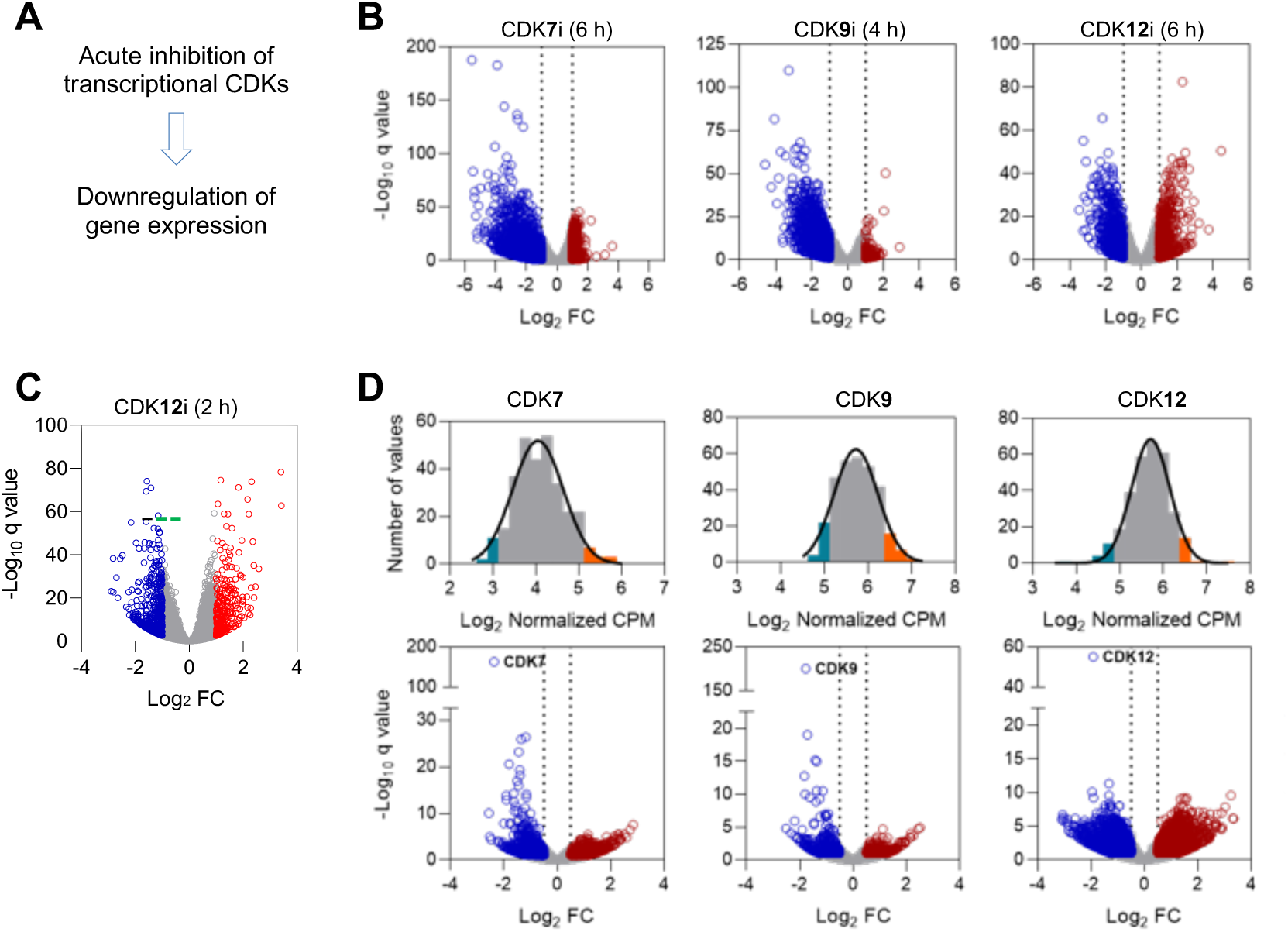
Gene upregulation is uniquely induced by CDK12 inhibition and is associated with low CDK12 expression in tumor samples. A. In theory, acute inhibition of transcriptional CDKs, particularly those involved in phosphorylating RPB1 CTD Ser2 phosphorylation, is expected to globally suppress gene expression, predominantly affecting mRNAs with short half-lives. B. Volcano plots of gene expression derived from 4-6 hours treatment of CDK7 inhibitor THZ1 (250 nM) in ovarian cancer cell line Kuramochi^9^, CDK9 inhibitor HH1 (10 µM) in a cell line (YB5) derived from the SW48 colon cancer cell line^10^, or CDK12 inhibitor SR-4835 (90 nM) in triple-negative breast cancer line MDA-MB-231^11^. RNA-seq data were downloaded from the Gene Expression Omnibus (GEO) and analyzed. C. A volcano plot of nascent RNA expression from neuroblastoma cells (IMR32) treated with 400 nM THZ531 for 2 hours^15^. Note that the sequencing involved 4-thiouridine-pulse labeling and included RNA spike-in control. D. (Top) selection of TCGA ovarian serous adenocarcinoma samples with low or high expression of the indicated CDK genes (the top and bottom 5% in terms of CDK7, CDK9 or CDK12 mRNA expression in ovarian cancer samples with expression data; n = 17 each group). (Bottom) volcano plots of differential gene expression in tumors with low expressing CDKs compared to those with high expressing CDKs. For all volcano plots, genes significantly upregulated or downregulated (absolute log_2_ fold change (FC) ≥ 1, p < 0.1) are colored in red and blue, respectively.

## Results

### Dual modes of gene expression alteration by CDK12 inhibition

To understand better the impact on gene expression of abolishing CTD phosphorylation, we first analyzed RNA-sequencing (RNA-seq) data based on acute inhibition of CTD kinases CDK7, CDK9 and CDK12. Consistent with the notion that acute inhibition of CTD kinases suppresses gene expression, administering the CDK7 inhibitor THZ1 for 6 hours in ovarian cancer cells^9^, or treating with the CDK9 inhibitor HH1 for 4 hours in colon cancer cells^10^ resulted in marked downregulation of gene expression (Figure 1B). However, inhibiting CDK12^11^, another transcriptional CDK that was also shown to phosphorylate CTD Ser2^11–14^, elicits a unique, dual pattern of alterations in gene expression (Figure 1B). In particular, gene upregulation following CDK12 inhibition is as pronounced as gene downregulation, indicating that the observed upregulation is likely not a secondary effect involving the de-repression of genes linked to downregulation. In addition, this robust upregulation of gene expression is also evident when nascent mRNAs were measured after neuroblastoma cells were exposed a CDK12 inhibitor for 2 hours^15^ (Figure 1C). Therefore, we sought to understand how CDK12 inhibition induces the expression of certain genes, when the global CTD Ser2 phosphorylation is reduced.

To confirm the observed dual effects on gene expression in other experimental systems, we examined RNA-seq data from studies utilizing genetic approaches to interfere with CDK12 activity^11, 16–18^. Remarkably, CRISPR or RNAi-mediated knockdown of CDK12, or inducible silencing of a Cdk12 transgene in embryonic stem cells with disrupted endogenous *Cdk12* alleles, all elicited both up- and down-regulation of gene expression (Figures S1A-S1D). Additionally, inducible CRSIPR/Cas9-mediated editing of Cyclin K, the cyclin partner of CDK12^14, 19^, led to a widespread upregulation of gene expression (Figure S1B). By nature, these genetic approaches do not interfere with CDK12 activity as rapidly as small molecule inhibitors, suggesting that the ensuing changes in gene expression are likely a combination of both primary and secondary effects of CDK12 loss. However, unlike CDK12 knockdown, CRISPR-mediated knockdown of CDK13^11^ – a transcriptional CDK that is closely related to CDK12 and also binds to Cyclin K, mainly causes gene downregulation (Figure S1A). Taken together, these findings suggest that genetic manipulation of CDK12 activity, similar to its acute inhibition by kinase inhibitors, triggers the upregulation of numerous genes and results in a dual-mode alteration of gene expression.

We next investigated whether the dual effects on gene expression also occur in human tumor samples. To this end, we analyzed tumors with low expression of CDK12 (CDK12_low) in comparison to those with high expression of CDK12 (CDK12_high). We reasoned that such an analysis requires that the two groups of samples have gene expression patterns lacking other heterogeneities that would confound differences in genome-wide expression. By plotting t-distributed stochastic neighbor embeddings (t-SNE) based on the normalized expression of genes associated with intrinsic subtyping^20–22^, we found that CDK12_low and CDK12_high samples within the TCGA Ovarian Serous Adenocarcinoma cohort, unlike datasets from breast or prostate cancer, demonstrate a reasonably homogenous pattern of gene expression (Figure S2A). We thus compared the gene expression of CDK12_low ovarian tumors with that of their CDK12_high counterparts (Figure 1D top). As a control, we performed similar analyses for tumors with low or high expression of CDK7 or CDK9 (Figure 1D top). Interestingly, tumors with low expression of CDK7 or CDK9 are associated with more significant downregulation than upregulation of gene expression (Figure 1D bottom). In contrast, CDK12_low tumors demonstrate differences in gene expression that are relatively greater in both magnitude and number but, more importantly, reflect an apparent dual-mode pattern in which gene upregulation is as significant as gene downregulation (Figure 1D, bottom right). We also compared the gene expression of 17 ovarian tumors with CDK12-inactivating alterations to that of an equally sized group of tumors with high CDK12 expression. Similar to ovarian tumors with low CDK12 expression, tumors with CDK12 mutations display pronounced differences in gene expression occurring in both directions (Figure S2B). These observations distinguish CDK12 from CDK7 or CDK9. Notably, it is unexpected that significant gene upregulation occurs following inactivation of a kinase traditionally considered to positively regulate transcription.

### CDK12 expression is associated with the progression of HER2+ breast cancer

This unique characteristic of CDK12 inhibition prompted us to investigate whether CDK12 represents a promising cancer target whose inhibition may impede cancer cell growth through novel mechanisms. Although inactivating mutations of CDK12 have been documented in cases of ovarian and prostate tumors^23–25^, CDK12 is known for its co-amplification with HER2/ERBB2 in HER2+ breast cancer^26, 27^ (Figure 2A and 2B). This subtype of cancer is diagnosed on the basis of HER2/ERBB2 amplification and overexpression and is managed with HER2-targeted therapies. Interestingly, co-amplification of genes in the HER2/ERBB2 amplicon does not necessarily translate into gene overexpression; for example, the IKZF3 gene is invariably amplified with HER2 but is not robustly overexpressed in HER2+ breast cancer (Figures S3A and S3B). To determine whether genes in the HER2/ERBB2 amplicon might be relevant to disease pathogenesis, we analyzed the relationship between gene expression and the histologic grade of disease – a clinical indicator of malignant progression. Notably, among all genes in the amplicon, CDK12 stands out as the gene most significantly correlated with disease severity, with its expression strongly associated with the histologic grade of HER2+ breast cancer (Figures 2C and S3C). In addition, CDK12 is found to be overexpressed in HER2+ breast cancer, as determined by both RNA-seq (Figures S3B and S3D) and proteomic profiling (Figure S3E). These observations suggest that CDK12 may represent a promising therapeutic target for HER2+ breast cancer.

**Figure 2.**
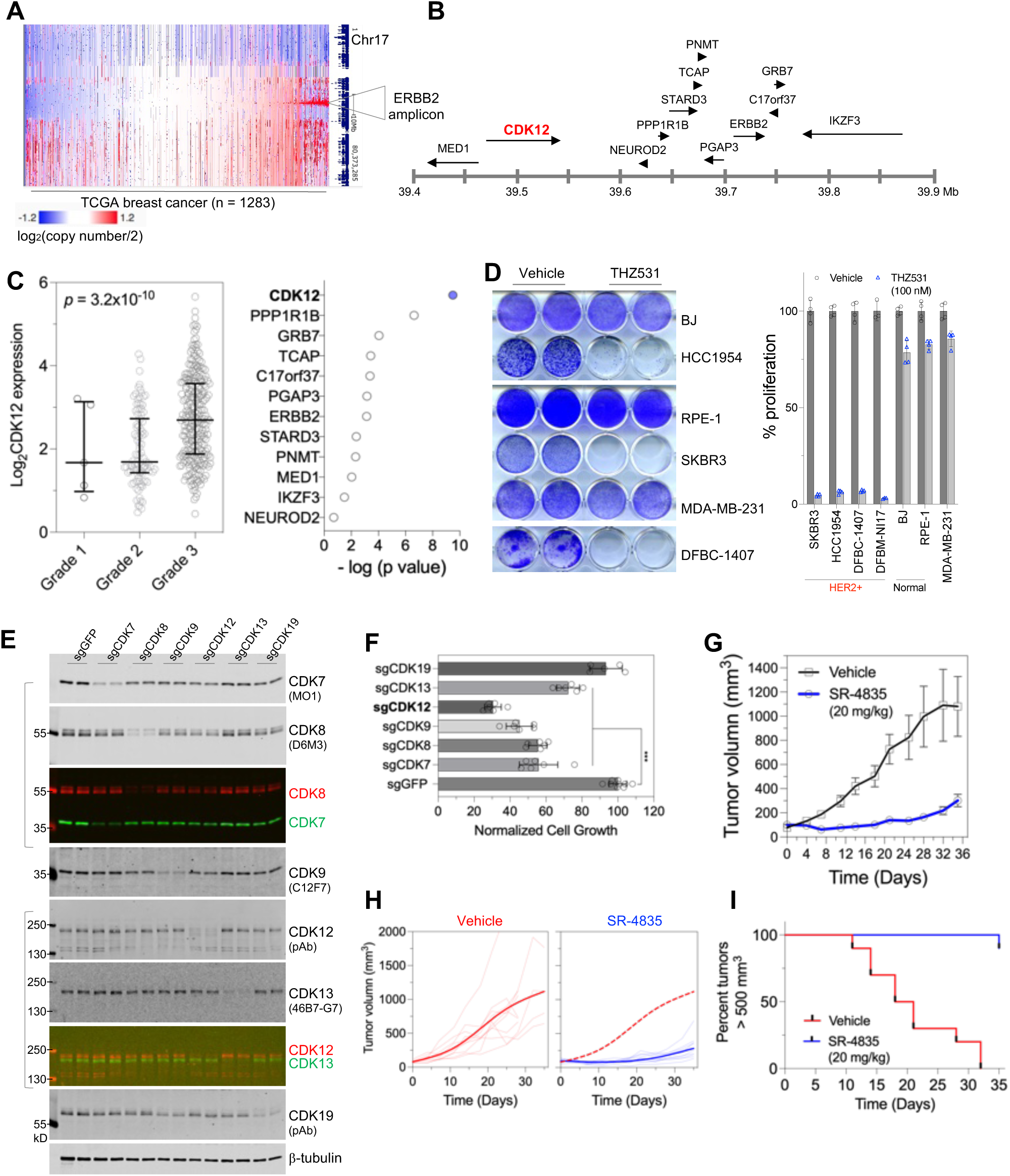
CDK12 inhibition impairs the growth of HER2+ breast cancer. A. Gene copy number status of human chromosome 17 (GDC TCGA breast cancer dataset). The heatmap was generated using the UCSC Xena Functional Genomics Browser (xena.ucsc.edu). Note that the HER2/ERBB2 amplicon occurs in approximately 15% breast cancer patients. B. A diagram depicting the structure of the HER2/ERBB2 amplicon in human chromosome 17. C. Association between the expression of HER2/ERBB2 amplicon genes and tumor grade. Correlation analysis was performed on gene expression in relation to the histologic grade of HER2+ breast cancer of the METABRIC cohort^60^, with p values generated from One-way ANOVA analysis. (Left) Plot for CDK12 gene expression in HER2+ tumors of different histological grades. (Right) A summary of the statistical significance for the association. D. The indicated cells or primary culture were treated with vehicle control or THZ531 (100 nM) for seven days, followed by fixation and crystal violet staining (left), and cell growth quantification (right). Note that DFBM-NI17 cells, which grow in suspension, had their cell growth measured in 96-well plates using CellTiter-Glo assays. E. HER2+ breast cancer cells (SKBR3) were transduced with lentiCRISPR virus targeting the indicated genes. After puromycin selection, cells were harvested for fluorescent immunoblotting. The molecular weights of the fluorescent protein markers and clone identities for monoclonal antibodies are indicated. Merged images show signals from two antibodies raised in different species. F. Cells as in (D) were seeded in 12-well plates at a density of 5,000 cells per well. After one week of incubation, cells were fixed and stained with crystal violet. The staining was subsequently solubilized for quantification of cell growth. ***p < 0.001. G. Mice with orthotopic xenografts of HER2+ breast cancer cells (HCC1954) were treated with SR-4835 (20 mg/kg by gavage) or vehicle (30% hp-BCD). Tumor size is shown as mean ± SE. H. Tumor size measurement of individual HCC1954 tumors over time (fine lines) and fitted tumor volume estimates (bold lines). Dashed red line in the right plot represents the fitted tumor growth curve from mice treated with vehicle. I. Kaplan-Meier curve demonstrating the percentage of animals with xenograft HER2+ tumors (HCC1954) smaller than 500 mm^3^ at the indicated time (p < 0.0001, log-rank [Mantel-Cox] test).

### Interfering with CDK12 activity impairs the growth and survival of HER2+ breast cancer cells

We proceeded to use small molecule inhibitors and CRISPR/Cas9-mediated gene editing to evaluate the function of CDK12 in HER2+ breast cancer. We first treated HER2+ breast cancer cell lines with THZ531, a covalent CDK12 kinase inhibitor^28^. Although HER2+ breast cancer cell lines displayed a variable response to the HER2 kinase inhibitor lapatinib, they all demonstrated exceptional sensitivity to THZ531, with half-maximal inhibition of cell growth at nanomolar concentrations (Figure S4A). THZ531 treatment induced apoptotic cell death, as indicated by appearance of cell fragmentation and PARP cleavage (Figures S4B and S4C).

Given the current challenges in treating relapsed HER2+ breast cancer in the clinic, including tumor metastasis and acquired resistance to multiple lines of HER2-targeted therapies^29, 30^, we derived primary cultures of HER2+ metastatic breast tumors and investigated their response to CDK12 inhibition. Different from HER2+ breast cancer cell lines that were established prior to the advent of cancer targeted therapies, the primary cultures were derived from metastasized tumors in patients who had received HER2-targeted therapies. Compared with HER2+ cancer cell lines, the primary cultures of HER2+ metastatic breast cancer were resistant to T-DM1 (Figures S4D and S4E), an antibody-drug conjugate approved for the treatment of HER2-positive advanced breast cancer^31^. Notably, these primary cultures of HER2+ metastatic breast cancer, similar to HER2+ breast cancer cell lines, were exquisitely sensitive to THZ531 (Figure S4F). Furthermore, HER2+ cancer cells demonstrated a markedly elevated sensitivity to CDK12 inhibition, as compared with untransformed human cells (Figure 2D).

To further study the role of CDK12 in HER2+ breast cancer cells, we knocked down the expression of CDK12 and examined its effect on cancer cell proliferation. In order to minimize the chances for counterselection, we subjected freshly infected and antibiotic-selected cells to clonogenic growth assays. CRISPR/Cas9-mediated CDK12 gene editing decreased CDK12 protein abundance, and impaired the clonogenic cell growth of HER2+ breast cancer cells. Notably, compared with other transcriptional CDKs including CDK7, CDK8, CDK9, CDK13, and CDK19, CDK12 gene editing elicited pronounced cell growth inhibition in HER2+ breast cancer cells (Figures 2E, 2F, S4G and S4H).

We next studied the anti-tumor potency of CDK12 inhibition in animal model of HER2+ breast cancer. In a xenograft model based on the HER2+ breast cancer cell line HCC1954, we administered an orally available CDK12 inhibitor, SR-4835^11^. Compared to vehicle control treatment, CDK12 inhibition caused a pronounced suppression of tumor growth, with the average tumor volume in SR-4835-treated mice less than one third of that in vehicle-treated group (Figures 2G-2I), while animal weight was not apparently altered^11^. Together, these data suggest that CDK12 represents a promising target for HER2+ breast cancer, an aggressive type of malignancy for which the demand for new drug targets is high due to the inevitable development of drug resistance to HER2-targeted therapies.

### Inhibiting CDK12 elicits dual modes of gene expression alteration in HER2+ breast cancer cells

We next examined CTD phosphorylation and gene expression in HER2+ cancer cells treated for a short period of time with CDK12 inhibitor THZ531, using CDK7 or CDK9 inhibitors as comparisons. Acute exposure to all three inhibitors efficiently decreased CTD Ser2 phosphorylation, as detected by both monoclonal and polyclonal antibodies recognizing this modification (Figures 3A, 4A and S5C Presumably). Presumably due to a more substantial reduction of RPB1 CTD phosphorylation, both CDK7 and CDK9 inhibition resulted in greater electrophoretic mobility of total RPB1 protein than CDK12 inhibition (Figures 3A, 4A and S5C). Therefore, THZ531-induced loss of CTD Ser2 phosphorylation signal is substantial, however, its impact on the overall phosphorylation status of RPB1 appears significantly smaller than that resulting from CDK7 or CDK9 inhibition.

**Figure 3.**
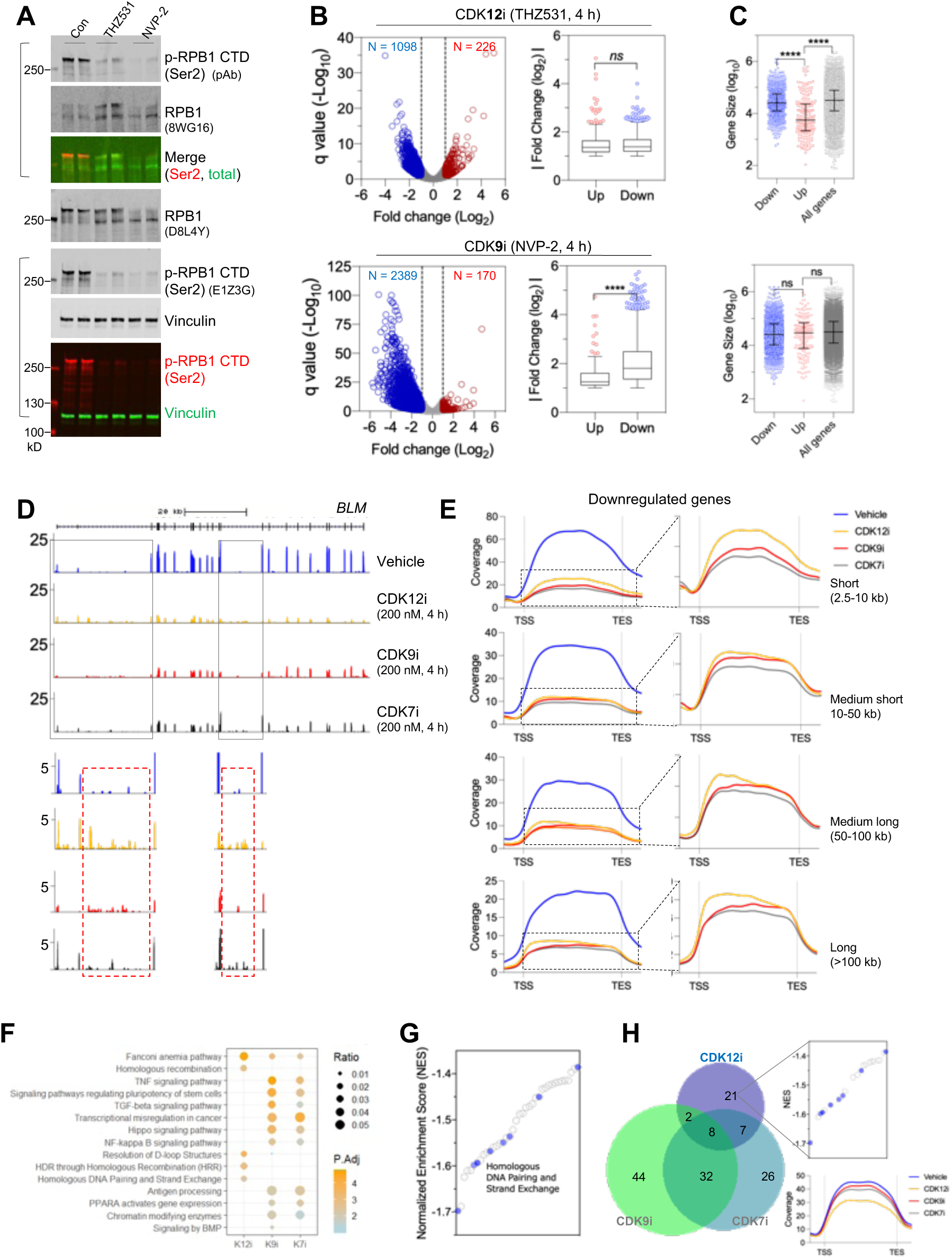
Dual modes of alteration of gene expression by CDK12 inhibition. A. HCC1954 cells were treated with THZ531 or NVP-2 for 4 hours, after which whole cell lysates were harvested for fluorescent immunoblotting. The molecular weights of the fluorescent protein markers and clone identities for monoclonal antibodies are indicated. Merged images show signals from two primary antibodies raised in different species. B. Volcano plots of gene expression. HCC1954 cells were treated for 4 h with THZ531 (200 nM), or NVP-2 (200 nM), or vehicle control. Total RNA was extracted and subjected to library construction and deep sequencing. Dotted lines in the volcano plots (left) indicate the thresholds used for log-transformed fold change (1, -1). The Tukey box plots (right) illustrate the comparison between the magnitudes of change in significantly up- or down-regulated genes. **** p < 0.0001; *ns,* not significant (Mann-Whitney test). C. Analysis of gene size among groups of genes impacted by CDK12 or 9 inhibition. **** p < 0.0001; *ns,* not significant (Mann-Whitney test). D. Traces of RNA-seq coverage over the gene BLM. Note that CDK12 inhibition decreases coverage over exons, but concomitantly produces reads at positions known to harbor intronic polyadenylation. The boxed regions are also displayed with re-scaled y-axis (bottom). E. Metagene profile plots of genes that are commonly downregulated in HCC1954 cells treated with THZ531, NVP-2, or THZ1 and that have been segmented by gene size. Note that CDK12 inhibition is uniquely associated with a size-dependent elongation defect. F. Overrepresentation analysis of significantly downregulated genes in HCC1954 cells that were treated as indicated in the x-axis. Gene sets in the KEGG and Reactome pathway databases were evaluated. Dot color denotes the statistical significance of gene set enrichment within the group of downregulated genes (-log q) and dot size denotes the fraction of downregulated genes corresponding to that gene set. G. Summary plot of normalized enrichment scores (NES) for gene sets that are significantly downregulated in HCC1954 cells treated with CDK12 inhibitor (nominal p-value < 0.05). Blue dots denote gene sets implicated in the DNA damage response. H. Venn diagram for gene sets that are downregulated by CDK12, CDK9, and CDK7 inhibition. Note that gene sets involved in the DNA damage response are specific to CDK12 inhibition. The right bottom illustrates metagene profile plots of genes in the gene set Reactome_ Homologous DNA Pairing and Strand Exchange in HCC1954 cells treated with THZ531, NVP-2, or THZ1 and that have been segmented by gene size.

**Figure 4.**
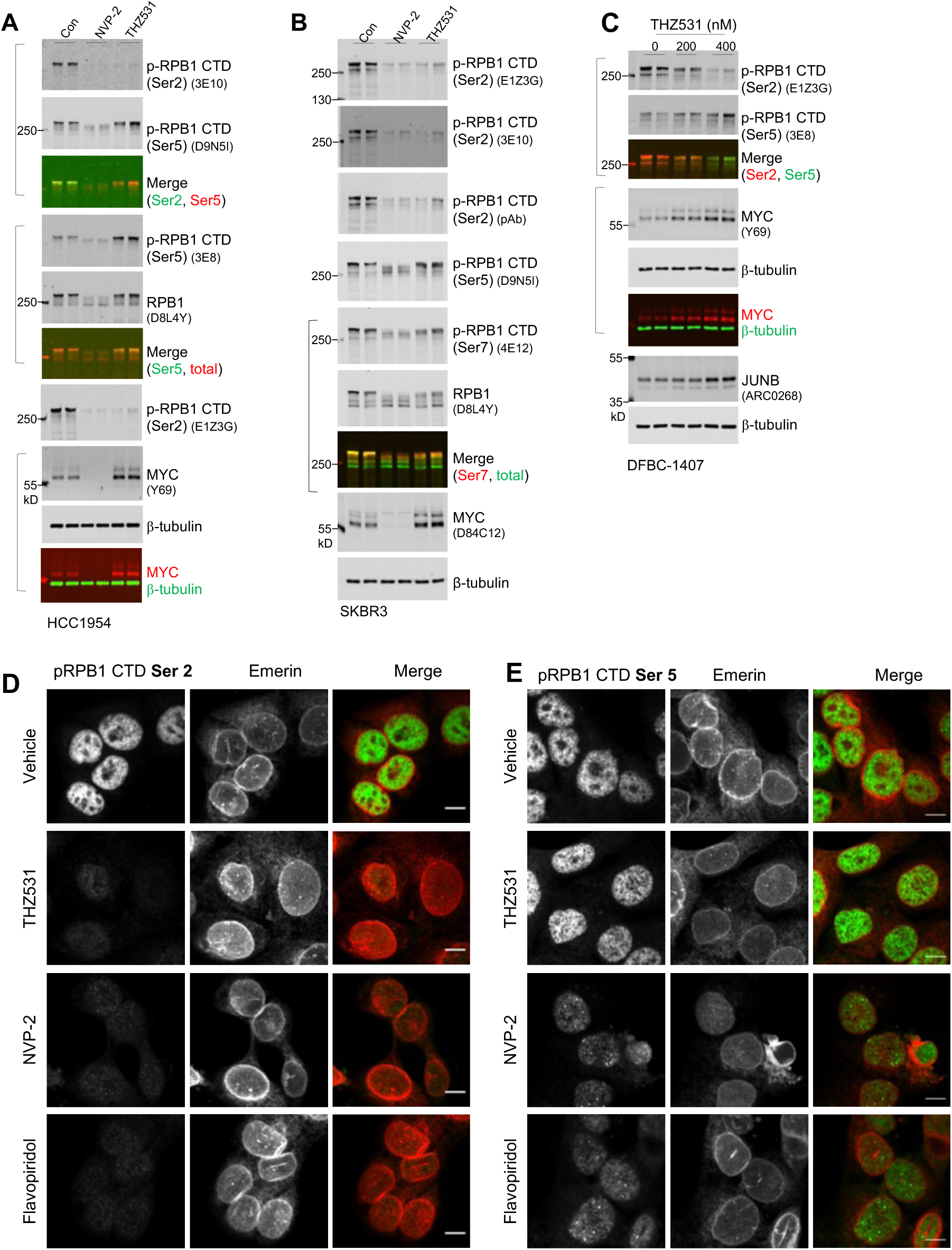
Enhanced CTD Ser5 phosphorylation upon CDK12 inhibition. A. Fluorescent immunoblotting of whole cell lysates. HER2+ cells HCC1954 were treated for 4 hours with vehicle, 200 nM THZ531, or NVP2. Following the treatment, cells were lysed with 1x SDS sample buffer to generate whole cell lysates for fluorescent immunoblotting. Each lane was loaded with lysates representing the same number of cells. The molecular weights of the fluorescent protein markers and clone identities for monoclonal antibodies are indicated. Merged images show signals from two primary antibodies raised in different species. B. HER2+ breast cancer cells (SKBR3) were treated as in (A), with total cell lysates subjected to fluorescent immunoblotting. C. The primary culture of HER2+ metastatic breast cancer DFBC-1407 were treated with THZ531 at the indicated concentrations for 4 hours, followed by total cell lysate preparation and fluorescent immunoblotting. D. Fluorescent images of cells stained with anti-phospho-RPB1 CTD (Ser2) (clone E1Z3G). HCC1954 cells growing on glass coverslips were treated with vehicle (0.8% DMSO, v/v), 400 nM THZ531, 200 nM NVP2, or 400 nM flavopiridol, fixed with 4% paraformaldehyde, and subjected to staining using the rabbit monoclonal anti-phospho-RPB1 CTD (Ser2) antibody (clone E1Z3G) together with the mouse monoclonal anti-Emerin antibody (clone CL0203). Images were acquired using identical parameters including exposure time and laser intensity on a spinning disk confocal microscope. The images displayed have the identical minimum and maximum displayed values among different treatment groups. Scale bar is 10 micrometers. E. Fluorescent staining for phospho-RPB1 CTD (Ser5) (clone D9N5I) alongside Emerin for nuclear membrane visualization. Cells were treated and stained as in (C). Confocal images were acquired using a 60x objective lens. Note that phospho-RPB1 (Ser5) signal persists in cells treated with THZ531, but diminished in cells exposed to NVP2 or flavopiridol.

Since CTD Ser2 phosphorylation is associated with transcriptional pause release and elongation, we profiled gene expression in cells treated with these inhibitor for 4 hours. Reminiscent of observations from data generated in other cancer cells (Figures 1B and 1C), acute inhibition of CDK12 elicited widespread alterations in gene expression (Figure 3B top left and S5A). Importantly, upregulated genes exhibited changes that are comparable in magnitude to their downregulated counterparts (Figures 3B top right and S5B left). In striking contrast to CDK12 inhibition, both CDK7 and CDK9 inhibition led to robust suppression of gene expression with much less potent induction of gene expression (Figures 3B bottom and S5C-S5E). These data indicate that in HER2+ cancer cells, similar to other systems (Figures 1B, 1C, and S1A-S1D), CDK12 inhibition evokes a dual-mode alteration in gene expression. For the groups of genes that were induced or repressed by THZ531, we found that upregulated genes are significantly shorter than either downregulated genes or all expressed genes in total (Figures 3C top and S5B right). Such a pattern appears to be unique to CDK12 inhibition and was not observed in cells treated with CDK7 or CDK9 inhibitors (Figures 3C bottom and S5E). Thus, consistent with prior studies^14, 15, 32–34^, our data indicate that transcriptional regulation by CDK12 has drastically different outcomes in short versus long genes.

Previous studies demonstrated that short-term inhibition or depletion of CDK12 increases the usage of intronic polyadenylation sites that are expected to give rise to truncated isoforms of long genes^15, 16^. Consistent with these findings, CDK12 inhibition in HER2+ cancer cells resulted in increased reads at locations known to trigger intronic polyadenylation (Figure 3D), and reduced the abundance of proteins encoded by these long genes (Figure S5F). We also found that CDK12 inhibition preferentially increased the usage of known proximal alternative polyadenylation sites in 3’ UTR (Figures S5G and S5H). Notably, CDK12 inhibition was distinctly associated with a size-dependent elongation defect in downregulated genes; this elongation defect results in elevated coverage towards the 5’ end of a gene (Figure 3E), a phenomenon that cannot be entirely explained by an induction of premature polyadenylation^15, 16^. Instead, these observations suggest that CDK12 inhibition potentially elicits defects in Pol II elongation that are increasingly manifested over the course of transcription.

Among the several hundred genes whose expression is altered by acute CDK12 inhibition, we investigated which gene(s) might represent key mediator(s) of the anti-proliferative capacity of CDK12 inhibition. To this end, we utilized gene set enrichment analysis (GSEA)^35, 36^ to identify gene sets sensitive to the intervention. Consistent with previous findings^11, 14–16, 28^, DNA damage response (DDR) genes – such as those catalogued in the signature of Homologous DNA Pairing and Strand Exchange – are significantly enriched among the downregulated genes (Figures 3F, 3G, and S6A-S6C) and were confirmed to be downregulated by quantitative PCR (Figures 6C and S8B). We also assessed gene set enrichment following acute CDK7 or CDK9 inhibition. Interestingly, the gene sets downregulated by disrupting CDK7 or CDK9, but not CDK12, share significant overlap relating to transcriptional machinery and signaling (Figures 3F and S6A). Notably, DDR genes are not enriched among the genes downregulated by CDK7 or CDK9 inhibitors (Figure 3H). These findings indicate that CDK12 inhibition can selectively downregulate DDR genes, a characteristic that might be leveraged to sensitize tumors to PARP inhibition^37, 38^. They also suggest that the mechanistic actions of CDK12 in gene expression control are fundamentally different from those of CDK7 and CDK9.

### CDK12 inhibition leads to disruption of early elongation control

To probe how CDK12 inhibition produces dual effects on gene expression, we examined CTD Ser5 phosphorylation, a modification that is known to accumulate on promoter-proximally paused Pol II^2, 3^. Despite a reduction of CTD Ser2 phosphorylation following THZ531 treatment, CTD Ser5 phosphorylation was surprisingly increased, as determined by two independent monoclonal antibodies 3E8 and D9N5I (Figure 4A). A persistent signal of CTD Ser5 phosphorylation was also observed in other HER2+ cancer cells including cancer cell lines and a primary culture of HER2+ metastatic breast cancer (Figures 4B and 4C). Notably, although CTD Ser2 phosphorylation was abolished by both CDK12 and CDK9 inhibition (Figures 4A, 4B, and 4D), CTD Ser5 phosphorylation was enhanced by CDK12 inhibition, but was reduced by CDK9 inhibition (Figures 4A, 4B, and 4E). These differential effects on CTD Ser2 versus Ser5 phosphorylation likely explain why the electrophoretic mobility of total RPB1 protein was largely unchanged by THZ531 treatment (Figures 3A, 4A, 4B and S5C). Given that the CTD Ser5 phosphorylation is a readout for early elongation complexes, the increased signal upon CDK12 inhibition suggests a persistence of paused or immature Pol II associated with the chromatin.

To define how treatment with THZ531 alters Pol II elongation, we performed Precision Run-on (PRO)-seq. This technique maps actively engaged Pol II across the genome at single nucleotide resolution, allowing us to directly probe how loss of CDK12 activity influences the transcription process. Importantly, these experiments were performed using spike-in controls, to allow absolute quantification of signal across conditions. Quantification of PRO-seq reads across gene bodies in HER2+ breast cancer cells HCC1954 confirmed that, despite widespread gene downregulation upon 2 h treatment with THZ531, a subset of transcribed loci had increased Pol II density (Figure S7A).

Indeed, genes that were upregulated by THZ531 treatment in RNA-seq data (Figure 3B) showed significantly more PRO-seq signal, consistent with increased transcription activity (Figure 5A). Likewise, genes downregulated in RNA-seq had significantly less PRO-seq signal across gene bodies (Figure 5A). Interestingly, composite metagene plots of the PRO-seq signal revealed that THZ531 treated cells had reduced promoter-proximal Pol II, with increased Pol II in early gene bodies (Figure 5B). This distribution is suggestive of increased pause release following THZ531 treatment. To test this, we calculated the pausing index, which is the ratio of promoter-proximal versus gene body PRO-seq reads where lower values indicate a higher degree of pause release^39^. These data confirm a significant and broad shift towards lower pausing across genes in cells treated with THZ531 (Figure 5C).

**Figure 5.**
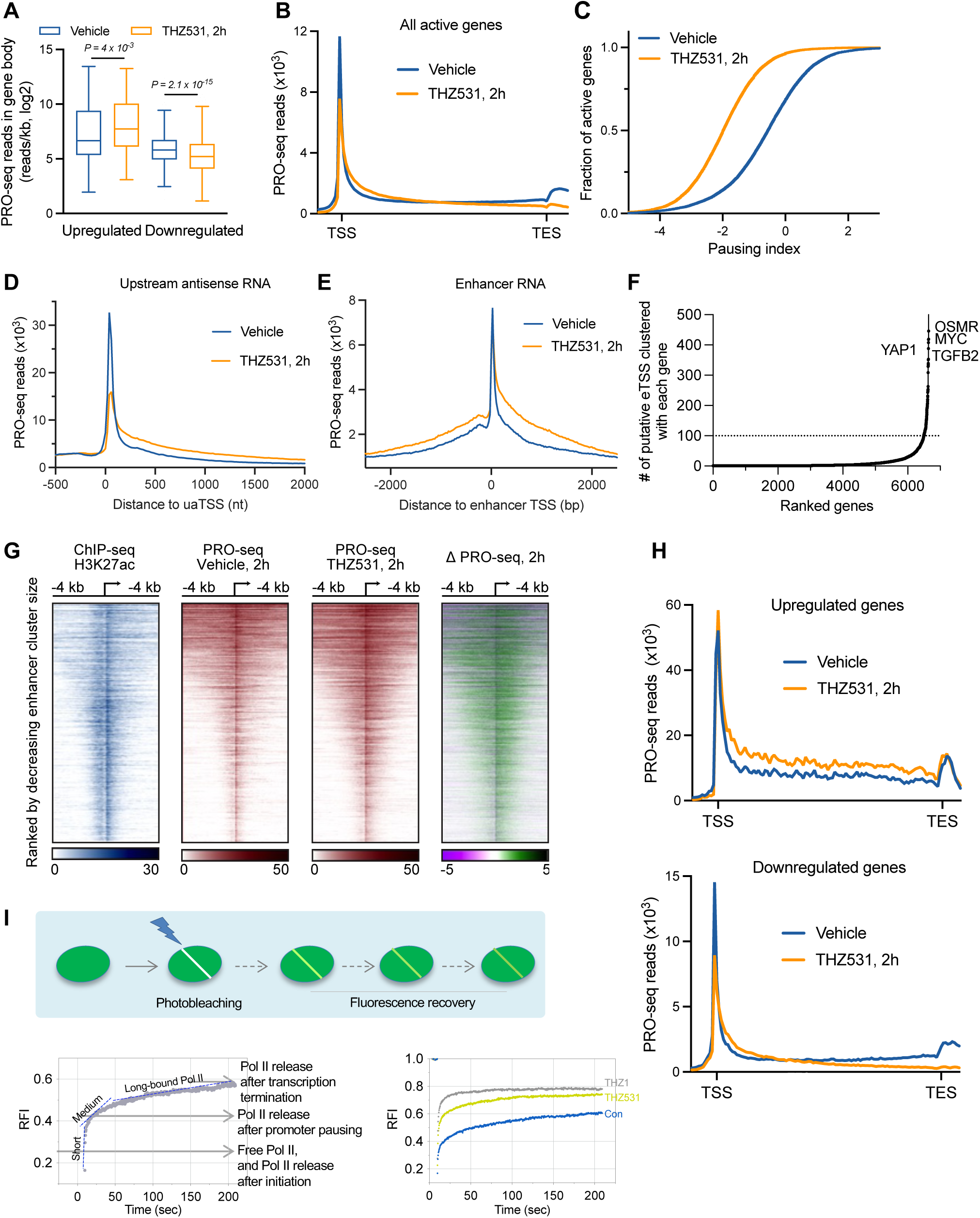
CDK12 inhibition leads to defective RNAPII elongation at coding and non-coding loci. A. Boxplots depict PRO-seq read density in mRNAs upregulated (N = 117) or downregulated (N = 805) upon THZ531 treatment, as defined in Figure 3B. Reads were counted from +250 nt downstream of the TSS to the transcript end site (TES), and normalized for gene length (reads/kb). Boxes show 25th–75th percentiles and whiskers depict 1.5 times the interquartile range. p values from Wilcoxon matched-pairs signed rank test. B. Metagene plot of average PRO-seq signal in vehicle and THZ531-treated HCC1954 cells across all active mRNA genes > 400 nt (N=11,876). Bins from TSS to TES are scaled to gene length, with 100 bins/gene, and data outside gene bodies are shown as average reads per gene in 200-nt bins. C. Cumulative distribution plot comparing the pausing indices of all genes shown in (B), comparing vehicle to THZ531 treatment. D-E. Metagene analysis of PRO-seq signal at (D) uaRNA loci (N = 9,617) or (E) eRNA loci (N=24,790) upon 2 h of indicated treatment. Data are shown as average reads per TSS in 25-nt bins. F. The distribution of the number of enhancers associated with the 13,530 active genes in HCC1954 cells. Association was based on the nearest active gene to each enhancer TSS. A subset of genes (n = 152; marked by dotted line) contains ≥ 100 associated eTSSs. G. Heat maps rank-ordered by decreasing size of each enhancer region. Shown are histone modifications (H3K27ac), and PRO-seq reads on both strands, with data centered around the dominant eTSS in each enhancer cluster (N = 7,631). H. Metagene plots of average PRO-seq signal in vehicle and THZ531-treated HCC1954 cells across genes defined as upregulated or downregulated in RNA-seq. Data are shown as in (B). I. Top cartoon: Measurements of Pol II diffusion kinetics determined by fluorescence recovery after photobleaching of a narrow strip spanning the nucleus (Strip-FRAP). (Bottom Left) FRAP was performed in untreated MRC-5 GFP-RPB1 KI cells and GFP-RPB1 fluorescence in the strip was background-corrected and normalized to pre-bleach fluorescence intensity and set at 1. The three dotted lines indicate kinetically distinct Pol II fractions. (Bottom Right) Pol II mobility in untreated cells or upon treatment with 400 nM THZ531 for 90 min, or 2 µM THZ1 for 90 min. Mean values from >13 nuclei of two independent experiments were plotted. RFI, Relative fluorescence intensity.

Strikingly, the upregulation observed upon THZ531 treatment was not limited to protein-coding genes, but extended across a broad range of RNA biotypes and genomic loci. Following THZ531 treatment, we detected a marked increase in PRO-seq reads downstream of upstream antisense RNA (uaRNA) start sites (Figure 5D), suggesting aberrant release of paused Pol II at these loci. Moreover, similar increases in PRO-seq signal were evident at enhancer regions throughout the genome (Figure 5E). Given that super-enhancers are known to drive robust gene expression^40, 41^, we further identified a subset of genes associated with large enhancer clusters (Figure 5F). Across all such super-enhancer clusters, THZ531 treatment led to a pronounced increase in PRO-seq signal, indicating a broad and global upregulation of enhancer activity (Figure 5G). These findings collectively demonstrate that THZ531 induces a pervasive transcriptional response, affecting both gene bodies and regulatory elements, and highlight the capacity of CDK12 inhibition to activate transcription at a genome-wide scale.

### Inhibition of CDK12 disrupts productive Pol II elongation of long genes, but allows synthesis across short genes

A widespread release of Pol II from promoter regions suggests either that pause release is increased by THZ531, or that the pausing checkpoint has been disrupted. Notably, when the early elongation checkpoint fails, the immature elongation complex released into genes is often defective in RNA synthesis^42, 43^. In agreement with this, our PRO-seq data suggests defective elongation across gene bodies (Figure 5B): despite more Pol II entering genes, we observe less Pol II reaching gene ends. As anticipated, this elongation defect was strongest at long genes, whereas short genes and non-coding RNAs exhibit increased PRO-seq signal across gene bodies (Figures S7B and S7C).

These findings suggest that genes upregulated in RNA-seq, which are significantly shorter than average (Figures 3C and S5B) would harbor engaged Pol II signal across the gene body in THZ531 treated cells, whereas the much longer, downregulated genes should show reduced PRO-seq signal towards gene ends. Analysis of PRO-seq data supports this model (Figure 5H), and emphasizes a striking reduction of Pol II at the end of downregulated genes.

Consistent with this notion, analyses of transient transcriptome sequencing (TT-seq) data^15^ show that, in response to THZ531, there was a global increase of reads near the transcription start site (TSS) (Figure S7D). However, the upregulated genes also exhibit a profound increase of reads extending as far as 5 kb away from TSS (Figure S7D), whereas downregulated genes begin to experience a rapid decrease of reads close to the 1 kb mark, indicating a reduction in productive elongation (Figure S7D).

We next studied the kinetics of Pol II diffusion in the nucleus following CDK12 inhibition, utilizing our previously developed model of GFP-RPB1 knock-in cells (Figure 5E top). Consistent with our previous studies^44^, we observed kinetically distinct Pol II fractions, representing initiating Pol II (designated short-bound Pol II), promoter-paused Pol II (medium-bound), and elongating Pol II (long-bound) (Figure 5E bottom left). As expected, THZ1 – a chemical known to block CTD phosphorylation and induce defects in co-transcriptional capping and pausing^45^ increased short-bound Pol II, coinciding with the loss of promoter-paused and elongating Pol II (Figure 5E bottom right). In contrast, CDK12 inhibition maintains the elongating Pol II fractions, as indicated by a fluorescence recovery that runs parallel to control group after 30 seconds, indicating that that the overall level of elongation is not reduced by THZ531 treatment. Together, these data suggest that CDK12 inhibition promotes immature elongation, and that the overall degree of elongation can remain largely unaltered due to the combined effects of enhanced immature elongation in early gene bodies and induced premature termination in long genes. Our data support previous studies indicating that THZ531 treatment inhibits productive elongation, reducing Pol II processivity^15, 32, 46^. Critically, our data also provide new insights into gene regulation following CDK12 inhibition, demonstrating aberrant pause release, which both contributes to the defects in elongation, and promotes increased synthesis of short genes.

### CDK12 inhibition activates MYC

Considering the unique role of CDK12 in regulating gene expression, we investigated whether gene activation contributes to the cancer cell growth inhibition conferred by CDK12 inactivation. Our analyses found that gene sets associated with transcriptional activity were enriched following CDK12 inhibition (Figure S6A), leading us to focus on a subset of upregulated genes encoding transcription factors. Intriguingly, MYC gene signatures were robustly activated by THZ531 (Figures 6A and S6B-6C), and by contrast, suppressed upon CDK7 or CDK9 inhibition (Figure S6D). Consistent with the observed activation of MYC gene signatures, THZ531 treatment led to a significant upregulation of MYC mRNA, as evidenced by mapping RNA-seq reads to its exons and confirmed by quantitative PCR (Figures 6B, 6C, S8A, and S8B). In HER2+ breast cancer cells including cell lines and a primary culture of HER2+ metastatic breast cancer, THZ531 treatment also increased MYC protein levels in a dose- and time-dependent manner, with 2-4 h treatments at 200-400 nM dosages resulting in pronounced increases in MYC protein (Figures 4A-4C, 6D, 6E, and S8C). Both the p64 and p67 isoforms of MYC^47^ were induced by CDK12 inhibition, with a similar fold change in induction detected by two different monoclonal antibodies recognizing MYC (Figure 6D). Notably, MFH-290, another selective CDK12 inhibitor with a chemical scaffold different from THZ531^48^, also increased MYC protein abundance in HER2+ breast cancer cells (Figure S8D). The MYC signals in the fluorescent immunoblots were from lysates of equal number of cells, indicating an induction of MYC protein *per* cell by CDK12 inhibition. In addition, the induced MYC protein depended on gene transcription and *de novo* protein synthesis, because it was abolished by RNA transcription inhibition with either the DNA intercalating anticancer drug actinomycin D or the Pol II inhibitor α-amanitin (Figures 6F and S8E), and by the protein synthesis inhibitor cycloheximide (Figures 6G and S8F).

**Figure 6.**
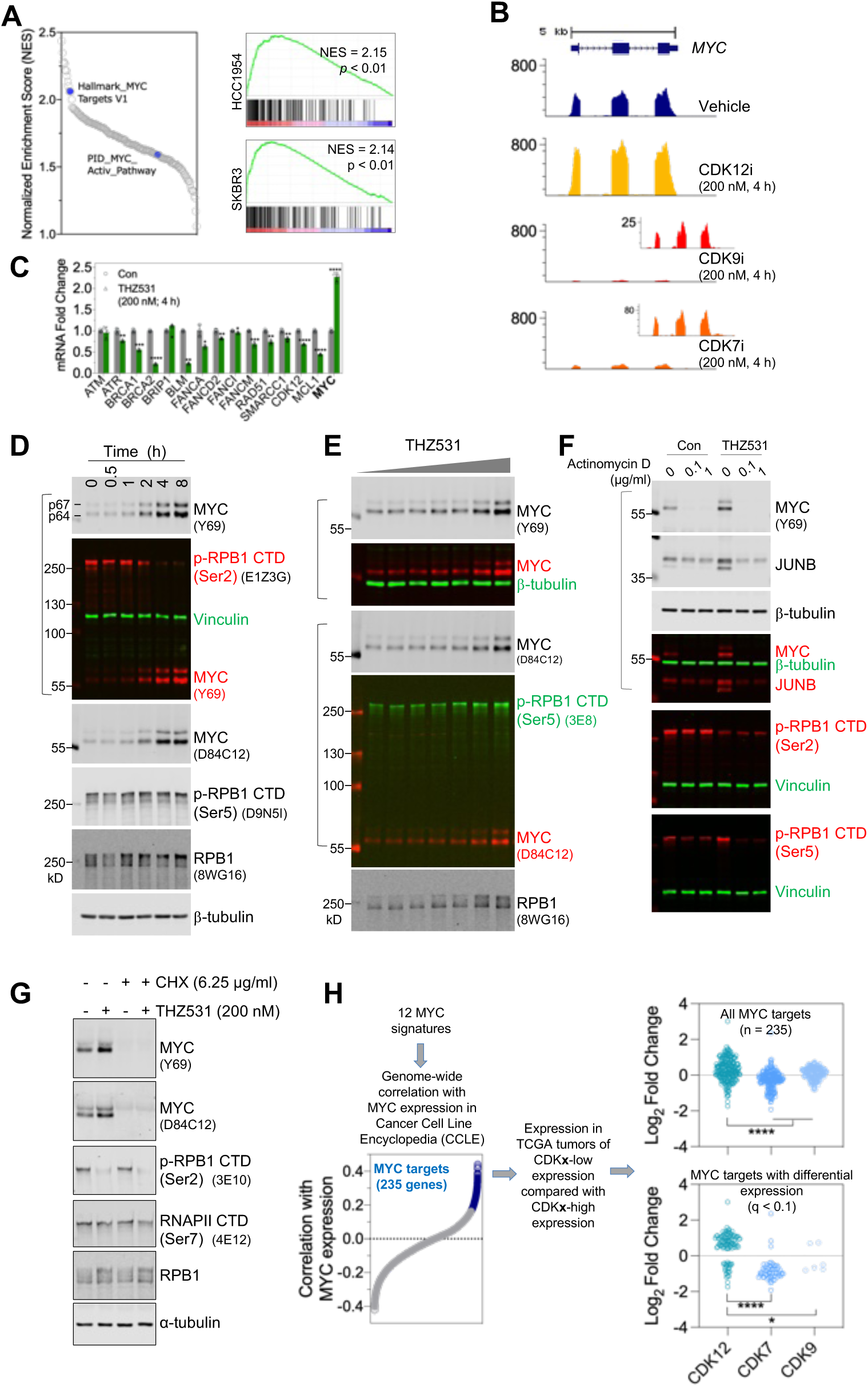
CDK12 Inhibition activates MYC. A. (Left) Summary plot of normalized enrichment scores (NES) for gene sets that are significantly upregulated in HCC1954 cells treated with CDK12 inhibitor (nominal p-value < 0.05). Blue dots denote MYC signature. (Right) GSEA plot of Hallmark_MYC Targets V1 for genes altered by THZ531 (200 nM, 4h) in HCC1954 (top) and SKBR3 (bottom) cells. Normalized enrichment score (NES) and p values are indicated. B. Traces of RNA-seq coverage over MYC in cells that were treated as indicated. Enlarged views for traces of CDK9 and CDK7 inhibition are shown as inserts with the y-axis scaled down. C. HCC1954 cells were treated as indicated for 4 h followed by total RNA extraction and reverse transcription. Quantitative PCR was then performed for the indicated genes. Note that MYC expression is increased, while other select DDR genes demonstrated significant downregulation. *p<0.05, **p<0.01, ***p<0.001 (Student’s t tests). D. HER2+ breast cancer cells were treated with THZ531 (200 nM) for the indicated time points. Cell lysates were prepared in SDS sample buffer and subjected to fluorescent immunoblotting. E. HCC1954 cells were treated with increasing concentrations of THZ531 (0, 12.5, 25, 50,100, 200, 400, and 800 nM) for 4 hours. Cell lysates were prepared and analyzed as in (G). F. HCC1954 cells were treated with Actinomycin D at the indicated doses in the presence of vehicle control or THZ531 (400 nM). Four hours post-treatment, cells were lysed with 1x SDS sample buffer, and lysates were prepared for fluorescent immunoblotting. Merged images are shown for blots using antibodies raised in different species. The very left lane was loaded with protein markers that emit near-infrared fluorescence. G. THZ531-induced MYC expression relies on de novo protein synthesis. HCC1954 cells were treated with cycloheximide (6.25 μg/ml) and THZ531 (200 nM), either individually or in combination. Cells were harvested in 3 hours for immunoblotting. H. Tumors with low expression of CDK12 exhibit activation of a MYC signature. (Right) The consolidated MYC signature was developed by combining all 12 gene sets from MSigDB (http://www.gsea-msigdb.org/) and identifying genes whose expression positively correlates with MYC across the 1305 cancer cell lines in the Cancer Cell Line Encyclopedia (CCLE, Pearson correlation *r* > 0.2). (Left) Expression of the 235 genes was then evaluated in CDK12, CDK7, CDK9-low ovarian tumors in comparison to their high expression counterparts. The top dot plot displays the differential expression of all 235 genes, while the bottom plot shows genes with significant differential expression (q < 0.1; n = 106, 60, 6 for the CDK12, CDK7, CDK9 groups, respectively). *p<0.05, ****p<0.0001 (Mann-Whitney test).

The profound induction of MYC protein appears to be unique to CDK12 inactivation, as inhibitors targeting CDK7 or CDK9, as well as the BRD4 inhibitor JQ1, resulted in a loss or reduction of MYC protein (Figures S9A-S9D). We consider this effect distinct from the compensatory induction of MYC following CDK9 inhibition, wherein MYC mRNA is first reduced after 2 h treatment but induced in the later timepoints^49^. Consistent with previous findings that inactivation of the mediator kinase CDK8/19 induces certain transcripts^50, 51^, MYC protein levels were modestly elevated in HER2+ breast cancer cells treated with the CDK8/19 inhibitor CCT251545^52^ (Figures S9A, S9B and S9E). Unlike HER2+ breast cancer cells, untransformed cells or triple-negative breast cancer (TNBC) cell lines did not have MYC induced upon CDK12 inhibition, despite the induction of dual-mode alteration of gene expression in TNBC cells (Figures 1C and S10C), and the induction of other immediate early genes, such as JUNB and FOS (Figures S10A and S10B). Nevertheless, MYC induction by THZ531 is not unique to HER2+ breast cancer cells and has been seen but not further studied in other types of cancer cells, such as neuroblastoma cells^15^, ovarian cancer cells^9^, BRAF-mutated melanoma cells^34^, and colon cancer cells^53^.

The findings in HER2+ cancer cells led us to investigate whether tumors with low CDK12 might exhibit enhanced MYC activity. To this end, we first refined MYC signatures by compiling all existing MYC signature gene sets deposited in the Molecular Signatures Database^54^ (MSigDB) and selecting those with expression that correlates significantly with MYC expression across the 1305 cancer cells lines in the Cancer Cell Line Encyclopedia^55^. We selected a total of 235 genes as the consolidated MYC signature and assessed their expression in ovarian tumors (Figure 6H left). Notably, ovarian tumors with low expression of CDK12 demonstrate higher expression of MYC signature genes (Figure 6H right). In contrast, tumors with low CDK7 expression exhibit lower expression of the MYC targets. Therefore, consistent with the observation that CDK12 inhibition activates MYC in HER2+ breast cancer cells, lower CDK12 activity in human tumors is associated with increased MYC expression and activity.

### A role for MYC induction in the antiproliferative potency of CDK12 inhibition

Widely known as a proto-oncogene, MYC exerts pleiotropic effects including the potent induction of apoptotic cell death upon forced activation in specific cells or tissues^56–58^. To determine whether MYC induction following CDK12 inactivation contributes to the observed cancer cell death and growth inhibition, we knocked down MYC expression in HER2+ breast cancer cells using CRISPR/Cas9-mediated gene editing (Figure 7A). sgMYC cells treated with THZ531 exhibited largely identical gene expression patterns compared with their treated parental cells (Figure 7B). However, they were distinct in their MYC status, with MYC signatures being significantly altered (Figures 7C, 7D, and S11A-S11E). Notably, sgMYC cells demonstrated increased resistance to CDK12 inhibition but were indistinguishable from control cells in response to the chemotherapeutic agent doxorubicin (Figures 7E and 7F), suggesting that the altered sensitivity to THZ531 is unlikely due to a possible change in drug efflux mechanisms. In addition, we generated a single clone of HER2+ cancer cells completely lacking both the p64 and p67 isoforms of MYC (Figure S11F). These MYC knockout cells also showed increased resistance to CDK12 inhibition (Figure S11G). Together, these data suggest that MYC acts as a key determinant of cellular response to CDK12 inactivation in HER2+ cancer cells.

**Figure 7.**
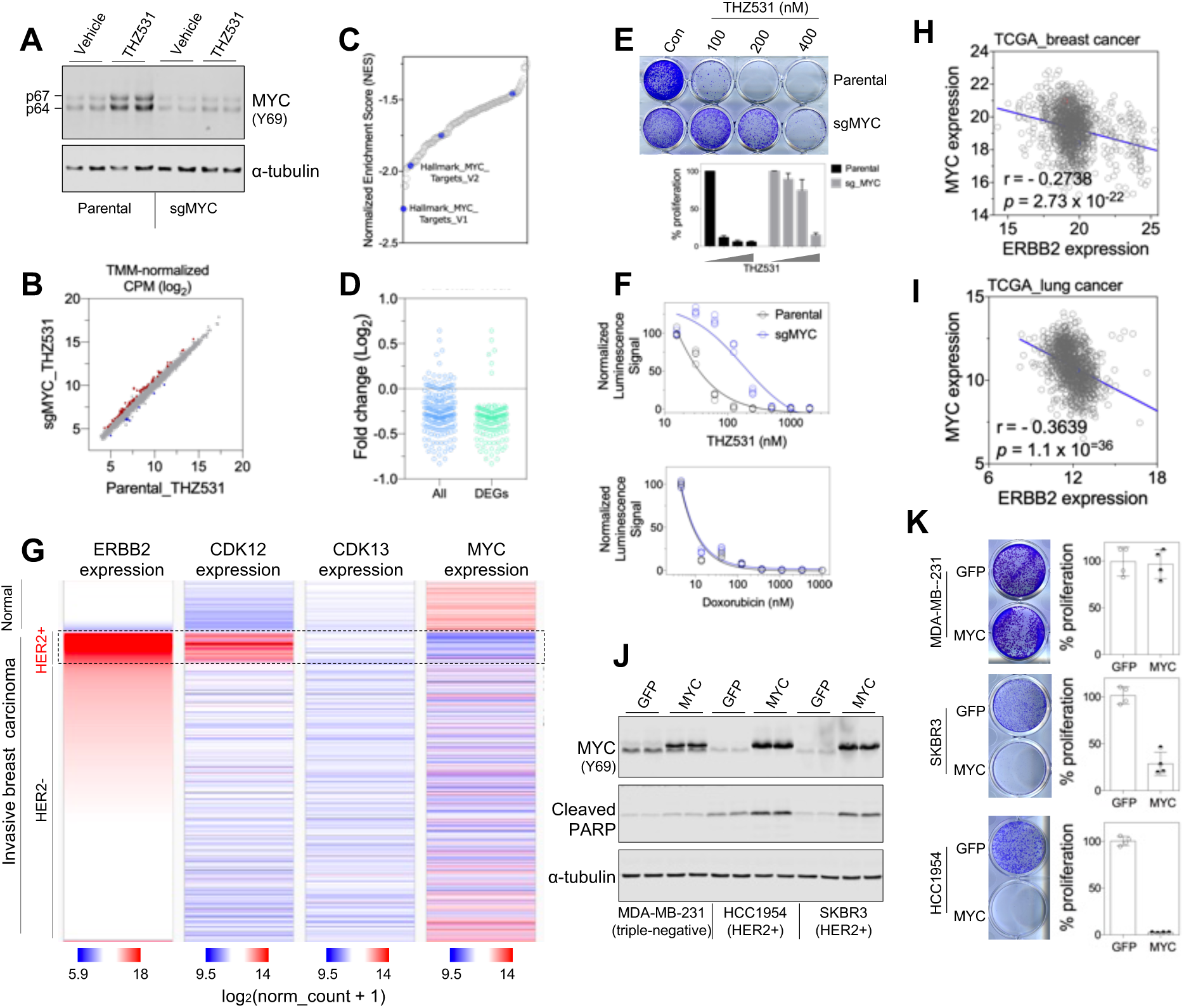
MYC mediates the anti-proliferative effect of CDK12 inhibition. A. MYC expression in parental HCC1954 cells and in HCC1954 cells following the infection of MYC-targeting lentiCRISPR vector (sg_MYC). The cells were treated with vehicle or THZ531 (200 nM) for 4 hours, followed by lysate preparation and fluorescent immunoblotting. B. Dot plot showing that treated sgMYC cells have similar global gene expression to treated parental cells. C. MYC signature is the top-ranked gene set among those downregulated in sgMYC cells treated with THZ531 relative to the treated, parental controls. Summary plot of normalized enrichment scores (NES) for gene sets that are significantly downregulated in sgMYC HCC1954 cells treated with CDK12 inhibitor (nominal p-value < 0.05). Blue dots denote MYC signatures. D. Expression of the consolidated 235-gene MYC signature in sgMYC HCC1954 cells treated with THZ531 compared to the treated, parental controls. The first column shows all genes, while the second shows the differentially regulated genes (DEGs, q<0.1). E. Response of control and sg_MYC cells to THZ531. The cells were treated with THZ531 at the indicated concentrations and, in 9 days, fixed for crystal violet staining (top) and subsequent quantification (bottom). F. CellTiter-Glo luminescent cell viability assays of parental and sgMYC cells treated with increasing concentrations of THZ531 (top) or doxorubicin (bottom) for 7 days. G. Heatmaps of ERBB2, CDK12, CDK13 and MYC expression in normal and malignant breast tissues. The data were generated with the use of UCSC Xena Functional Genomics Browser. Note that HER2+ breast tumors (highlighted) exhibit high expression of HER2/ERBB2 and CDK12, but lower expression of MYC compared to normal breast tissues. H. Plot of ERBB2 and MYC expression in breast tumor samples (TCGA). The Pearson correlation coefficient (*r*) and p value were calculated in GraphPad Prism 7. I. Reverse correlation between ERBB2 and MYC expression in lung cancer samples (TCGA). J. Fluorescent immunoblotting for analysis of MYC and cleaved PARP. Cells were infected with lentivirus, and harvested six days after the initial virus infection for the preparation of lysates. Note that apoptotic cell death, indicated by the appearance of cleaved PARP, occurs upon forced MYC overexpression in two HER2+ breast cancer cells, but is negligible in triple-negative MDA-MB-231 cells. K. Effect of MYC overexpression on the growth of cancer cells. The cells were transduced wit freshly packaged virus and, on day 5, seeded in 12-well plates (5,000 MDA-MB-231 cells, 20,000 HCC1954 or SKBR3 cells seeded in each well). Cells were fixed in 6 – 8 days upon reaching visual confluence and stained with crystal violet (left). The staining was then dissolved for the quantification of cell growth (right).

A potential role of MYC activation in mediating the killing of HER2+ cancer cell by CDK12 inhibition is surprising. Does MYC act as a tumor promoter or suppressor in human HER2+ breast cancer? Intriguingly, low MYC levels consistently co-occur with high HER2/ERBB2 expression. HER2+ breast cancers show the lowest MYC expression across all subtypes (Figure 7G), and a striking negative correlation between MYC and HER2/ERBB2 is observed in both TCGA^59^ and METABRIC^60^ datasets (Figures 7H and S12A). Since HER2/ERBB2 amplification and overexpression is not restricted to breast cancer^61, 62^, we further analyzed MYC status in malignancies of other tissue types and found a similar pattern: a statistically significant reverse correlation between MYC and HER2/ERBB2 expression exists across diverse cancer types, including lung cancer (r = -0.36; Figures 7I and S12B), bladder cancer (r = -0.41; Figure S12C), cervical cancer (r = -0.34; Figure S12D), kidney clear cell carcinoma (r = -0.47; Figure S12E), kidney papillary cell carcinoma (r = -0.43; Figure S12F), and prostate cancer (r = -0.23; Figure S12G). Furthermore, analysis of HER2+ breast cancers and normal breast tissues revealed a striking difference: normal breast tissues express significantly higher MYC levels while having lower HER2/ERBB2 and CDK12 expression as expected (Figures 7G and S12H). To circumvent confounding factors such as tumor heterogeneity in the analysis of bulk tumor RNA-seq data, we examined single-cell RNA sequencing data from human breast tumors^63^. Consistent with observations from bulk RNA-seq data analyses, tumor epithelial cells in HER2+ breast cancer exhibit the highest expression of HER2/ERBB2 and CDK12 but the lowest expression of MYC, compared with their counterparts in estrogen receptor positive (ER+) or triple-negative breast cancer (Figure S12I). Together, these observations suggest that MYC, though an established oncogene, is unlikely to be involved in promoting the development of HER2+ breast cancer. Instead, it appears that high HER2/ERBB2 activity in HER2+ breast cancer and other malignancies causes an inherent intolerance of MYC activation. To test this hypothesis, we transduced cells for overexpressing MYC and subjected the cells to growth assays right after transduction and antibiotics selection. Notably, in HER2+ but not in TNBC cells, overexpression of MYC impaired cell proliferation and triggered widespread apoptotic cell death as evidenced by induced PARP cleavage (Figures 7J and 7K). These findings support the notion that cancer cells with hyperactive HER2/ERBB2 signaling are inherently intolerant to MYC activation, providing a plausible explanation for the observed negative correlation between MYC and HER2/ERBB2 expression across diverse tumor types.

## Discussion

Leveraging small molecule inhibitors of transcriptional CDKs coupled with sequencing of nascent and steady-state RNA, we discovered that CDK12 inhibition reduces the expression of many genes but also simultaneously enhances the expression of other genes, especially those of small sizes. Our study points to a novel transcription mechanism – a key role of CDK12 in the pause release and early elongation checkpoints – to explain the co-occurrence of simultaneous gene down- and up-regulation. CDK12 inhibition interferes with the early elongation checkpoint and therefore causes a markedly increased but precocious Pol II release into gene bodies. The elongating Pol II generated by CDK12 inactivation tends to dissociate from the gene bodies earlier than normal. As a consequence, long genes will have incomplete transcription and instead premature termination and RNA decay, while short genes can still yield mature transcripts at increased levels. Such a dual-mode gene expression alteration upon CDK12 inhibition is fundamentally different from that resulting from CDK7 or CDK9 inhibition, which predominantly causes global gene downregulation. Our study proposes a CDK12-regulated elongation checkpoint that follows CDK7-controlled promoter clearance and CDK9-catalyzed pause release.

Although we studied altered gene expression following CDK12 inactivation in HER2+ breast cancer cells, the gene expression pattern regulated by CDK12 is not limited to the genetic background of HER2 or CDK12 gene amplification. We have analyzed a large body of sequencing data generated by independent studies using either CDK12 inhibition or genetic inactivation of CDK12 and found that the dual-mode gene expression control by CDK12 also exists in triple-negative breast cancer cells^11, 18^, neuroblastoma cells^15^, ovarian cancer cells^9^, murine leukemia cells^17^, and mouse embryonic stem cells^16^. Notably, our results are consistent with several recent studies reporting gene induction effects resulting from CDK12 inhibition, on short genes including those encoding protein components of the AP-1 and NF-κB pathways in BRAF-mutated melanoma cells^34^, and on genes driven by p53 and NF-κB in colon cancer cells^53^, but extend our understanding of CDK12 inhibition to non-coding loci like antisense promoter and enhancers. Therefore, the induction of gene expression by CDK12 inhibition appears to be a universal phenomenon in mammalian cells, while the identity of the induced genomic coding and non-coding loci may depend on the cellular context.

The model we propose to explain the dual-mode gene expression alteration elicited by CDK12 inactivation has three key elements. First, CDK12 is critical to maintain a Pol II early elongation checkpoint, as suggested by a marked increase of Pol II in the early regions of both down- and up-regulated genes upon acute CDK12 inhibition. Notably, these observations are found by PRO-seq studies in HER2+ breast cancer cells upon THZ531 treatment (Figures 5B-5D, S7C), as well as in other nascent RNA sequencing studies that we analyzed, including TT-seq data generated in neuroblastoma cells upon THZ531 treatment^15^ (Figure S7D). Second, the increased Pol II in early elongation regions is likely premature, because the signal decreases over time along the gene bodies, a phenomenon that is more apparent for long genes. Third, and as a consequence, long genes are exquisitely sensitive to CDK12 inhibition and their transcription more frequently ends with premature termination and decay.

The currently prevalent model of transcriptional regulation by CDK12 suggests that CDK12 inhibits intronic polyadenylation^15, 16^. In comparison, the model we propose for CDK12 regulation of an early elongation checkpoint can readily explain the reduction of CDK12-dependent long genes (*i.e.*, through premature termination and RNA decay), as well as the induction of short genes upon CDK12 inhibition. However, a role of CDK12 in checkpoint control of early elongation does not exclude its function(s) in directly regulating elongation and co-transcriptional splicing^33, 46, 64, 65^.

The effect of CDK12 inhibition in increasing early elongating Pol II and in inducing expression of short genes is reminiscent of what we recently found upon disrupting the Integrator complex^42, 43^. Different from CDK12 inhibition, cells with acute loss of Integrator subunits INTS1 or INTS11 have drastically less effect in reducing gene expression. This difference may result from the effect of CDK12 inhibition, but not INTS11 protein degradation, in reducing RPB1 CTD Ser2 phosphorylation - a modification known to be critical for the recruitment of elongating factors^66^. Therefore, the premature elongating Pol II in CDK12-inactivated cells may well be more impaired than Pol II in cells with acute inactivation of Integrator, leading to a lesser degree of gene induction while causing reduction of many long genes.

Our study finds that CDK12 inactivation induces the expression of MYC and MYC target genes, as part of the dual-mode alterations in gene expression regulated by CDK12. In HER2+ breast cancer cells, MYC induction results in cell death and growth inhibition, likely due to an inherent intolerance of HER2+ cells to MYC activation. Therefore, CDK12 could be a key therapeutic target in HER2+ breast cancer and, possibly, in other HER2+ malignancies where CDK12 is co-amplified and overexpressed. The therapeutic utility of CDK12 inhibition merits further preclinical and clinical investigation, given that HER2+ tumors inevitably develop resistance to multiple lines of HER2-targeted therapies. Although CDK12 acts as a proto-oncogene in HER2+ malignancies, it has inactivating mutations in other cancer types, such as in approximately 6% of castration-resistant prostate cancer^23, 25^. In addition, recent studies found that inactivation of Cdk12 in mouse prostate gives rise to preneoplastic lesions^67^. Whether CDK12 acts as tissue type-specific oncogenic factor or tumor suppressor in a given tumor type will be an important area of study.

In summary, our work reveals a key role of CDK12 in regulating an early transcriptional elongation checkpoint, with CDK12 inactivation inducing premature, but weakened, elongation into gene bodies. As a consequence, the precociously released Pol ll manifests elongation defects, leading to premature termination and RNA decay – an effect that is disproportionally apparent in long genes; in the meantime, many short genes have increased production of full-length transcripts suitable for protein synthesis. The dual-mode regulation of gene expression by CDK12 together with the magnitude of alteration is unique among transcriptional kinases and represents a novel, yet critical, module of gene expression control.

## ACKNOWLEDGEMENTS

We thank Drs. Richard Young (Whitehead Institute), Nathanael Gray (Stanford), Steve Buratowski (Harvard Medical School) for discussion and gifts of reagents. We thank the Core for Imaging Technology & Education (CITE) at Harvard Medical School for the use of confocal microscopes. We thank the HMS Nascent Transcriptomics Core for assistance with PRO-seq, and the Harvard Bauer Core Facility and HMS Biopolymers Facility for next-generation sequencing. We thank all members of the Adelman, Zhao and Roberts groups for helpful discussion and assistance throughout this project.

## Author contributions

All the authors designed the experiments and discussed the results. Y.W., C.M. and A.B. performed all the experiments and bioinformatic analysis, supervised by K.A., J.J.Z and T.M.R. J.T performed animal experiments, supervised by Y.W. J.N developed key reagents used in the study. R.J.J performed the kinetics analysis of Pol II diffusion, supervised by J.A.M. Y.W. wrote the manuscript with inputs from all authors.

## Data and Material availability

The RNA-seq and PRO-seq data generated in this study has been deposited in the Gene Expression Omnibus database (accession no. GSE181905, GSE302011). The following secure token has been created to allow review of record GSE302011 while it remains in private status: kjopqwyonjovxut.

---

**Figure S1.**
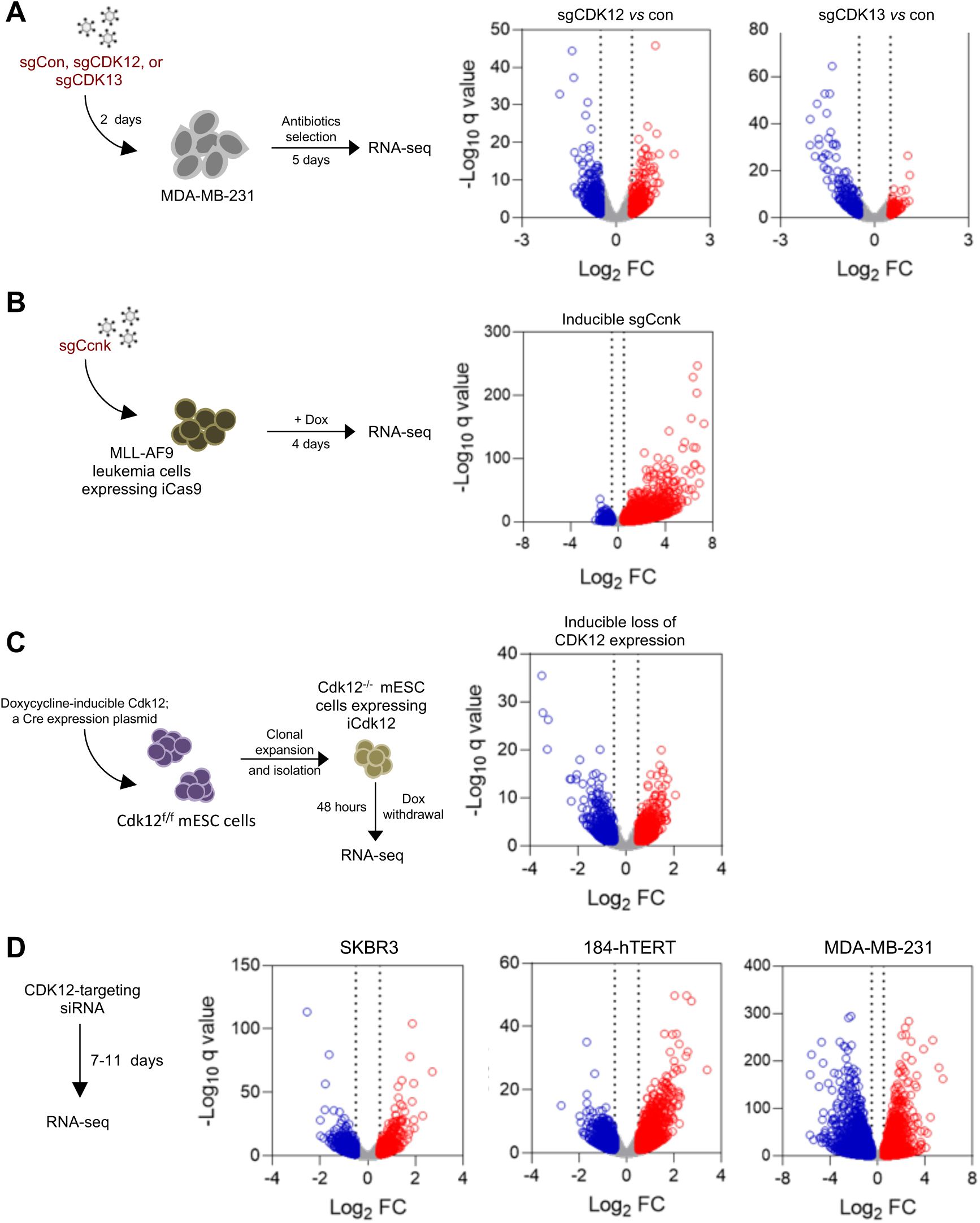
Loss of CDK12 expression causes a dual-mode alteration in global gene expression. A. Perturbing CDK12 in breast cancer cells by CRISPR-mediated gene editing^11^ results in gene upregulation as well as downregulation. In contrast, knocking down CDK13 primarily leads to gene downregulation B. Inducible knockdown of *Ccnk* in mouse leukemia cells^17^, the cyclin for CDK12 and CDK13, results in profound upregulation of gene expression. C. In mouse embryonic stem cells with ablated endogenous Cdk12 alleles, repressing the expression of transgenic *Cdk12*^16^ induces both downregulation and upregulation of gene expression. D. Silencing CDK12 expression with siRNA in HER2+ (SKBR3) or triple negative breast cancer cell line (MDA-MB-231), as well as a non-transformed breast cell line (184-hTERT)^18^, results in dual-modal alteration of global gene expression. All RNA-seq data were obtained from the NCBI Gene Expression Omnibus (GEO) and analyzed as described in the Methods section. Information regarding experimental procedures is from the original publications^11, 17, 18^. In the volcano plots, genes significantly upregulated or downregulated (absolute log_2_ fold change ≥ 1, p < 0.1) are colored in red and blue, respectively.

**Figure S2.**
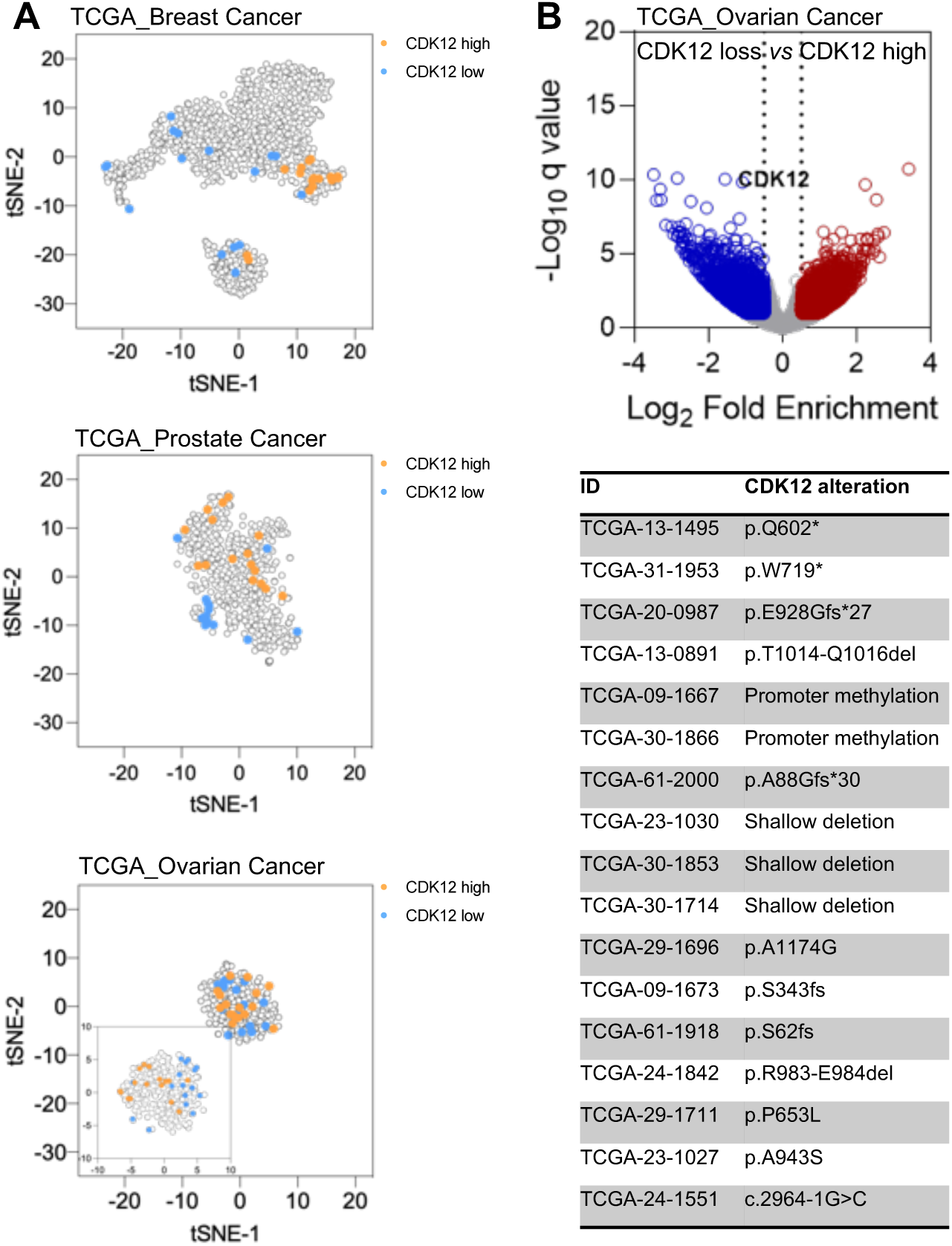
Comparison between tumors with loss of CDK12 expression and those with wild type CDK12. A. T-SNE plots for TCGA breast cancer (top), prostate caner (middle), and ovarian cancer (bottom) samples, each clustered according to PAM50 gene signature. All three plots are on the same x- and y-axis scales. The insert in the ovarian plot provides a zoomed-in view. Note that compared with breast cancer, ovarian cancer has a more homogenous expression of PAM50 genes, and that unlike prostate cancer, the CDK12 low or high ovarian samples are more dispersed out rather than clustered together. B. (top) A volcano plot comparing gene expression in ovarian tumors with CDK12 deletion or loss-of-function mutations to tumors with high expression of CDK12. (below) The table lists all TCGA ovarian tumors samples with CDK12 gene alterations.

**Figure S3.**
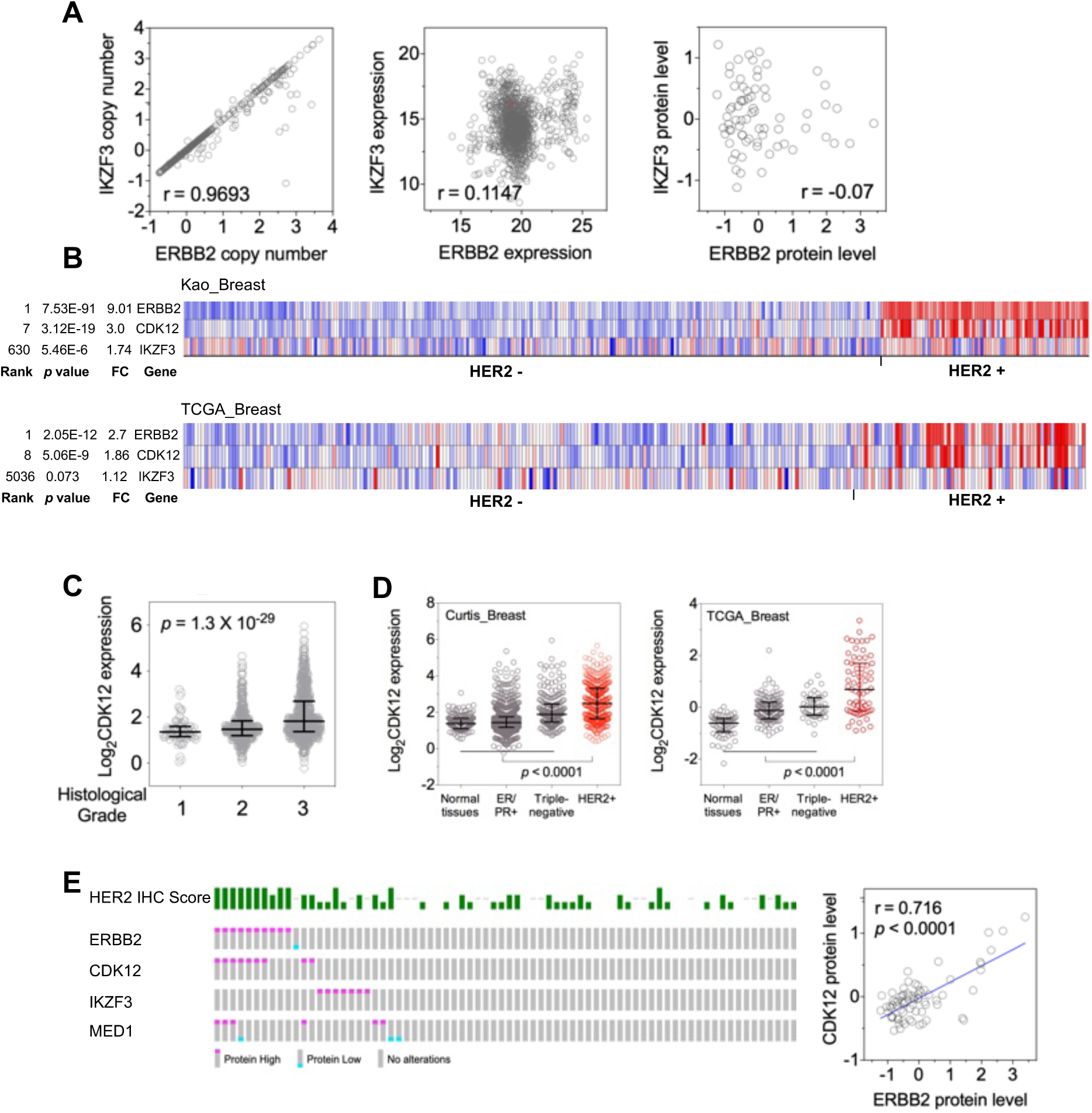
CDK12 gene expression among subtypes of breast cancer, and its association with disease status. A. An example showing that genes within the HER2/ERBB2 amplicon could have vastly different patterns of expression. The left dot plot indicates a nearly perfect correlation between the copy number of ERBB2 and that of IKZF3, the middle dot plot depicts a statistically weak correlation between the gene expression of ERBB2 and IKZF3, and the right dot pot shows a lack of correlation between the protein abundance of ERBB2 and IKZF3. Gene copy number and expression (RNA-seq) data in the GDC TCGA breast cancer cohort and proteomic data (Mertins et al., 2016) were analyzed in GraphPad Prism. The Pearson correlation coefficient (*r*) is indicated. B. Expression of indicated genes in HER2 negative (HER2-) or HER2 positive (HER2+) breast tumors. The analysis was performed via Oncomine (www.oncomine.org; Rhodes et al., 2007) for the cohorts of Kao_breast (Kao et al., 2011) and TCGA_breast (Cancer Genome Atlas Network et al., 2012). The statistical comparison between HER2+ and HER2-tumors include FC (fold change of gene expression), p value, and genome-wide ranking that is based on p values. C. Analysis of CDK12 gene expression in relation to the histologic grade of breast cancer in total cohort of breast cancer. CDK12 gene expression and the annotation of histologic grade of tumors were generated in the Curtis cohort (Curtis et al., 2012). The indicated *p* values were generated from One-way ANOVA analysis. D. Expression of CDK12 in normal breast tissues, ER/PR+, triple-negative, or HER2+ breast cancers. The expression data were generated in the Curtis (METABRIC)^60^ or TCGA cohort^59^. E. (Left) Abundance of indicated proteins in human breast cancer samples. The oncoprint visualization was derived from a query into the dataset of TCGA Breast Invasive Carcinoma at cBioPortal for Cancer Genomics (www.cbioportal.org; Gao et al., 2013). A total of 74 samples were subjected to mass-spectrometry-based proteomic analysis (Mertins et al., 2016). (Right) CDK12 protein abundance in breast cancer is significantly correlated with that of ERBB2. Quantification data were downloaded from cBioPortal and analyzed in GraphPad Prism 7. The Pearson correlation coefficient (*r*) is indicated.

**Figure S4.**
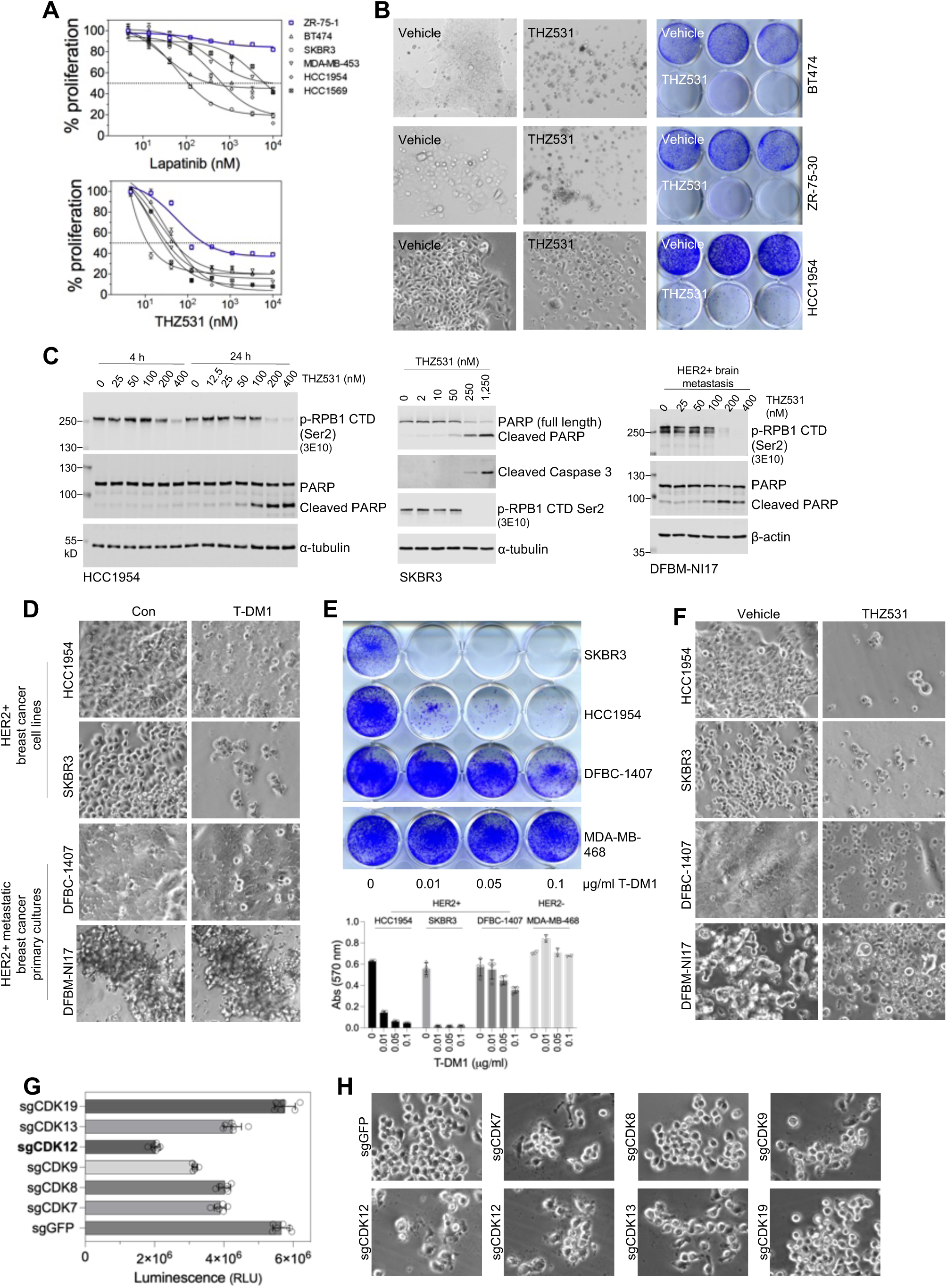
CDK12 inhibition suppresses cell growth and induces apoptotic cell death in HER2+ breast cancer cells. A. The anti-proliferative effects of the HER2 kinase inhibitor Lapatinib (top) and the CDK12 inhibitor THZ531 (bottom) in HER2+ breast cancer cell lines (black curves). Note that the ER+ breast cancer cell line ZR-75-1 (blue curve) showed resistance to Lapatinib while exhibiting modest sensitivity to THZ531. B. HER2+ breast cancer cells were seeded in 12-well plates and treated with vehicle control (DMSO) or THZ531 (50 nM for BT474 and ZR-75-30; 200 nM for HCC1954). Upon reaching visual confluency in the group of vehicle treatment, cells were imaged using a 10x objective lens (left). Cells were then fixed and stained with crystal violet. C. Induction of apoptotic cell death by THZ531. HER2+ breast cancer cells were treated with increasing concentrations of THZ531 for 24 h, unless specified otherwise. Cell lysates were prepared for fluorescent immunoblotting. The molecular weights of the fluorescent protein markers are indicated. D. Response of HER2+ breast cancer cells to T-DM1. The indicated HER2+ cancer cell lines and primary cultures of HER2+ metastatic breast cancer were exposed to T-DM1 (100 ng/ml for HCC1954, SKBR3, and DFBC-1407; 20 ng/ml for DFBM-NI17). Images were acquired after seven days of incubation using a 10x objective lens. E. Cells were treated as indicated, fixed, and stained with crystal violet (top). The staining was subsequently extracted for quantification of cell growth (bottom). Note that the triple-negative breast cancer cells (MDA-MB-468) were used as a negative control and demonstrated complete resistance to T-DM1. F. HER2+ breast cancer cell lines and primary cultures of HER2+ metastatic breast cancer were treated with vehicle control (0.02% DMSO, v/v) or THZ531 (100 nM). After seven days, bright-field images were captured using a 10x objective lens. G. HER2+ breast cancer cells (SKBR3) were transduced with lentiCRISPR virus targeting the indicated genes. After puromycin selection, cells were subjected to seven-day cell proliferation assays measured by CellTiter-Glo. Raw luminescence values of each group were plotted (mean ± SD). Unpaired Student’s t tests revealed significant differences between sgGFP and all sgCDK groups (p < 0.0001), except sgCDK19. Note that sgCDK12 cells demonstrated the most impaired cell proliferation. H. Bright-field images of SKBR3 cells transduced with indicated guides as in (G). Cells were seeded in 12-well plates at a density of 5,000 cells per well. After one week of incubation, images were captured using a 10x objective lens.

**Figure S5.**
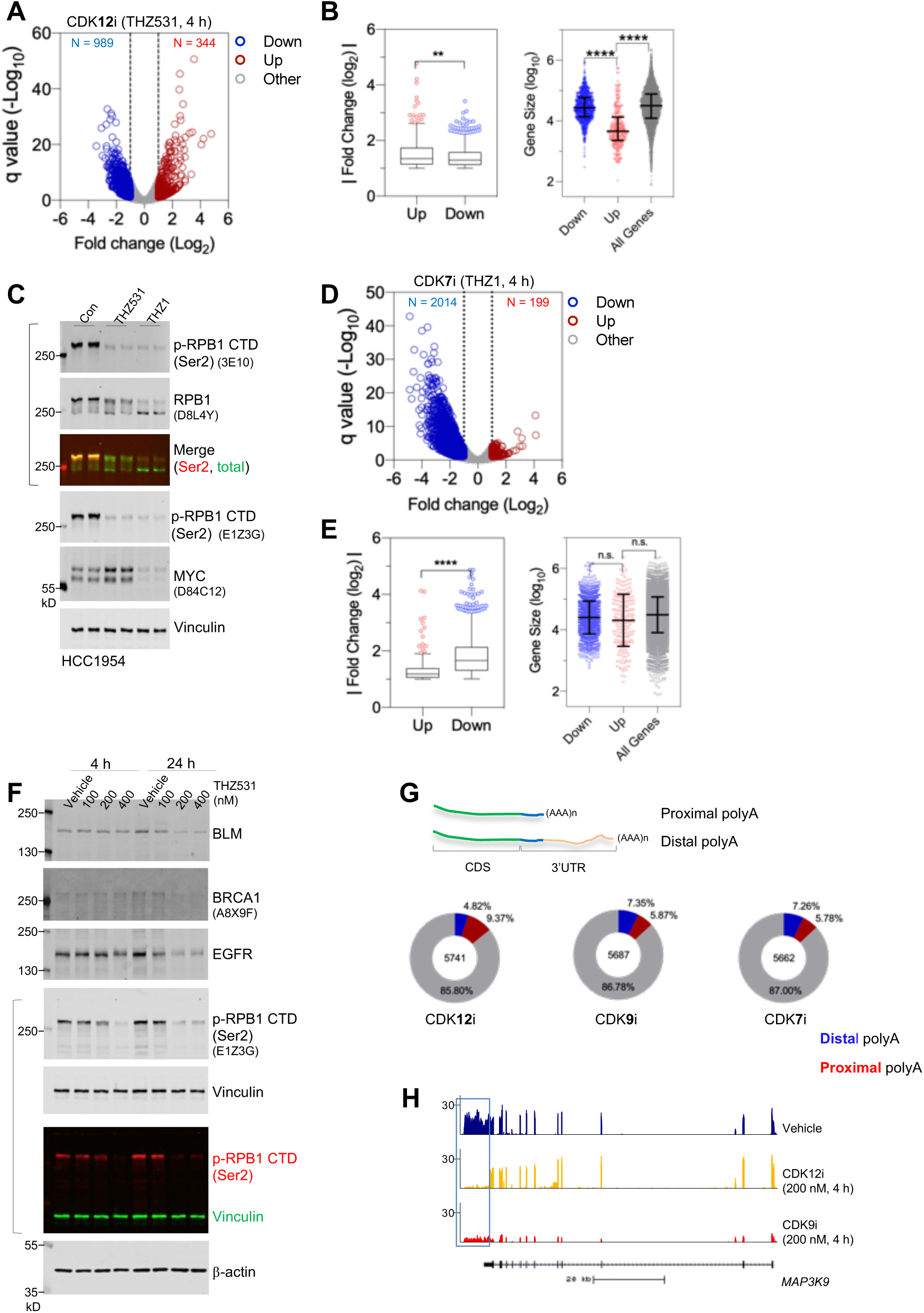
Differential impacts on gene expression by CDK12, CDK9, and CDK7 inhibition. A. A volcano plot of gene expression in SKBR3 cells treated with THZ531 (200 nM, 4 h) compared with vehicle control (0.04% DMSO, v/v). Blue and red dots denote individual genes with significant change in expression (fold of change > 1, q < 0.1). B. (Left) A Tukey box plot indicates that, for genes significantly altered in expression by THZ531 treatment in SKBR3 cells, the magnitude of upregulation is even greater than gene downregulation (**p < 0.01, Mann-Whitney test). (Right) Comparison of gene size among groups of genes that differentially respond to CDK12 inhibition (****p < 0.0001, Mann-Whitney test). C. HCC1954 cells were treated with vehicle control, 200 nM THZ531 or THZ1 for 4 h, followed by lysis with 1x SDS sample buffer and fluorescent immunoblotting. Note that THZ1 treatment caused a complete electrophoretic mobility shift of total RBP1, indicating a substantial loss of CTD phosphorylation. Note that THZ1 also targets CDK12 and CDK13, and thus lacks a desired selectivity. THZ1 was chosen for the current study, instead of a more selective version of CDK7 inhibitor YKL-5-124, primarily because YKL-5-125 does not have any effect on CTD phosphorylation (Olson et al., 2019). D. A volcano plot of gene expression in HCC1954 cells treated with THZ1 (200 nM, 4 h) compared with vehicle control. Blue and red dots are genes with significant change in expression (fold of change > 1, q < 0.1). E. (Left) A Tukey box plot indicating that, for genes with expression significantly altered by THZ1 treatment in HCC1954 cells, the magnitude of downregulation is greater than gene upregulation (****p < 0.01). (Right) A comparison of gene size among groups of genes that differentially respond to CDK12 inhibition (n.s., not significant; Mann-Whitney test). F. Fluorescent immunoblotting of total cell lysates from HCC1954 cells treated with indicated doses of THZ531 for 4 or 24 hours. Note that cells treated with THZ531 for 24 hours demonstrate a reduced protein abundance of ATM, BRCA1, and EGFR, all of which are encoded by large genes (>100 kb). G. (Top) A schematic depicting mRNA with proximal or distal polyA. (Bottom) The percentage of mRNA in each treatment group (HCC1954) showing positive (proximal polyA) or negative (distal polyA) change of proximal polyadenylation site usage (△PPAU). H. Traces of RNA Seq coverage over the MAP3K9 gene. Note that CDK12 inhibition in HCC1954 cells abolishes reads corresponding to a distal polyA site (boxed) while exerting no effect on coverage over the exons.

**Figure S6.**
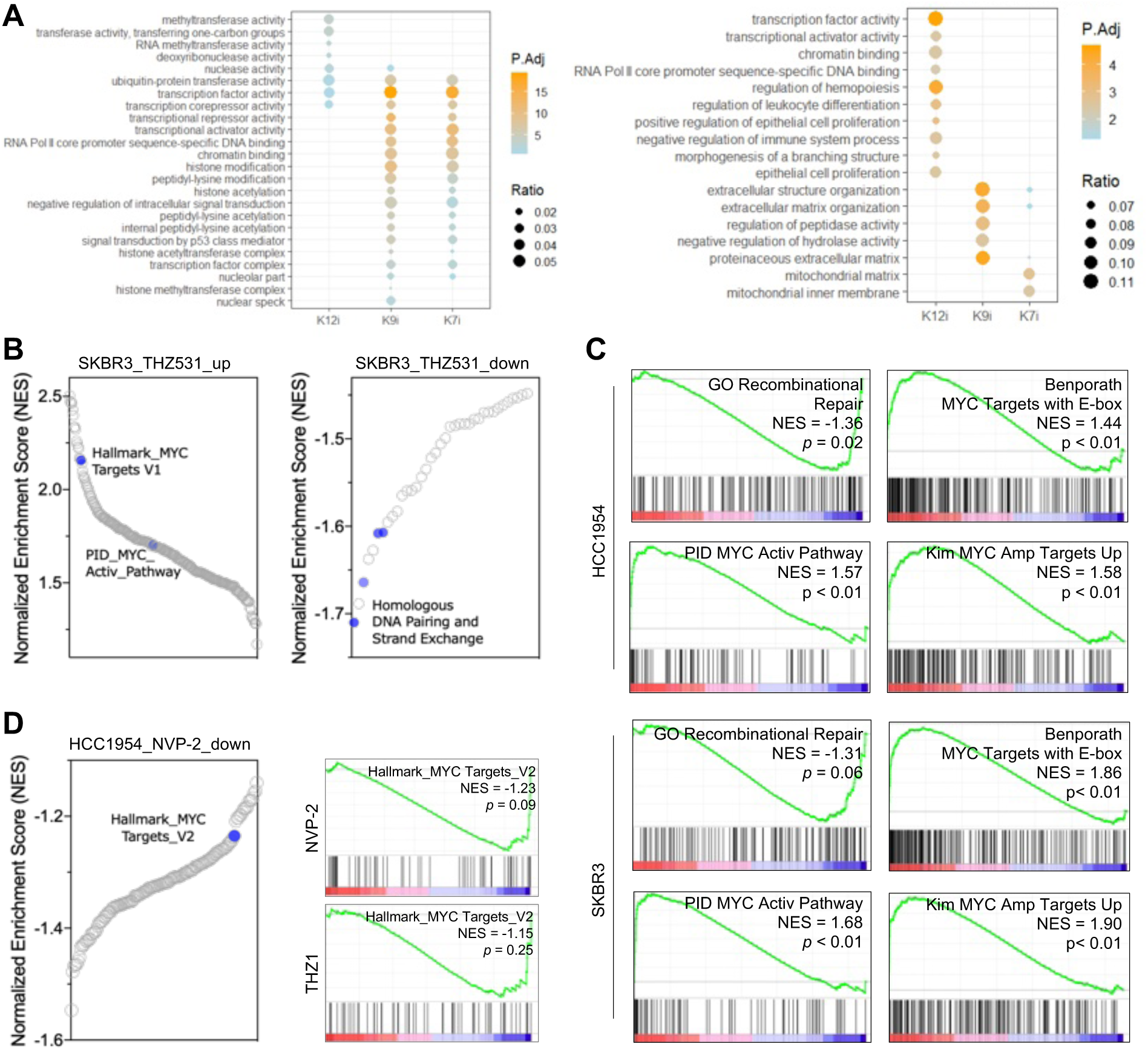
Acute CDK12 inhibition activates MYC. A. Overrepresentation analysis of significantly downregulated (left) or upregulated (right) genes in each of the treatments. Gene sets from the GO term database (> 10,000 gene sets) were evaluated. Dot color indicates the statistical significance (-log q) of the gene set representation within the group of down- or upregulated genes, and dot size represents the fraction of down- or upregulated genes corresponding to each gene set represented. B. Normalized enrichment score (NES) plots for gene sets that are significantly upregulated (left) or downregulated (right) in SKBR3 cells following CDK12 inhibition (nominal p-value< 0.05). Two MYC gene sets are highlighted and indicated (left), and 4 gene sets implicated in DNA damage response are highlighted in blue (right). C. GSEA plots for indicated gene sets in HCC1954 (top) and SKBR3 (bottom) cells treated with THZ531. NES and p values are indicated. D. NES plot for gene sets that are significantly downregulated in HCC1954 cells treated with the CDK9 inhibitor NVP-2 (nominal p-value < 0.05). The blue dots denotes MYC signature. The right plots depicts GSEA analysis of Hallmark_MYC Targets V2 gene set for genes downregulated by NVP-2 (200 nM, 4 h; top) and THZ1 (200 nM, 4 h; bottom). NES and p values are indicated.

**Figure S7.**
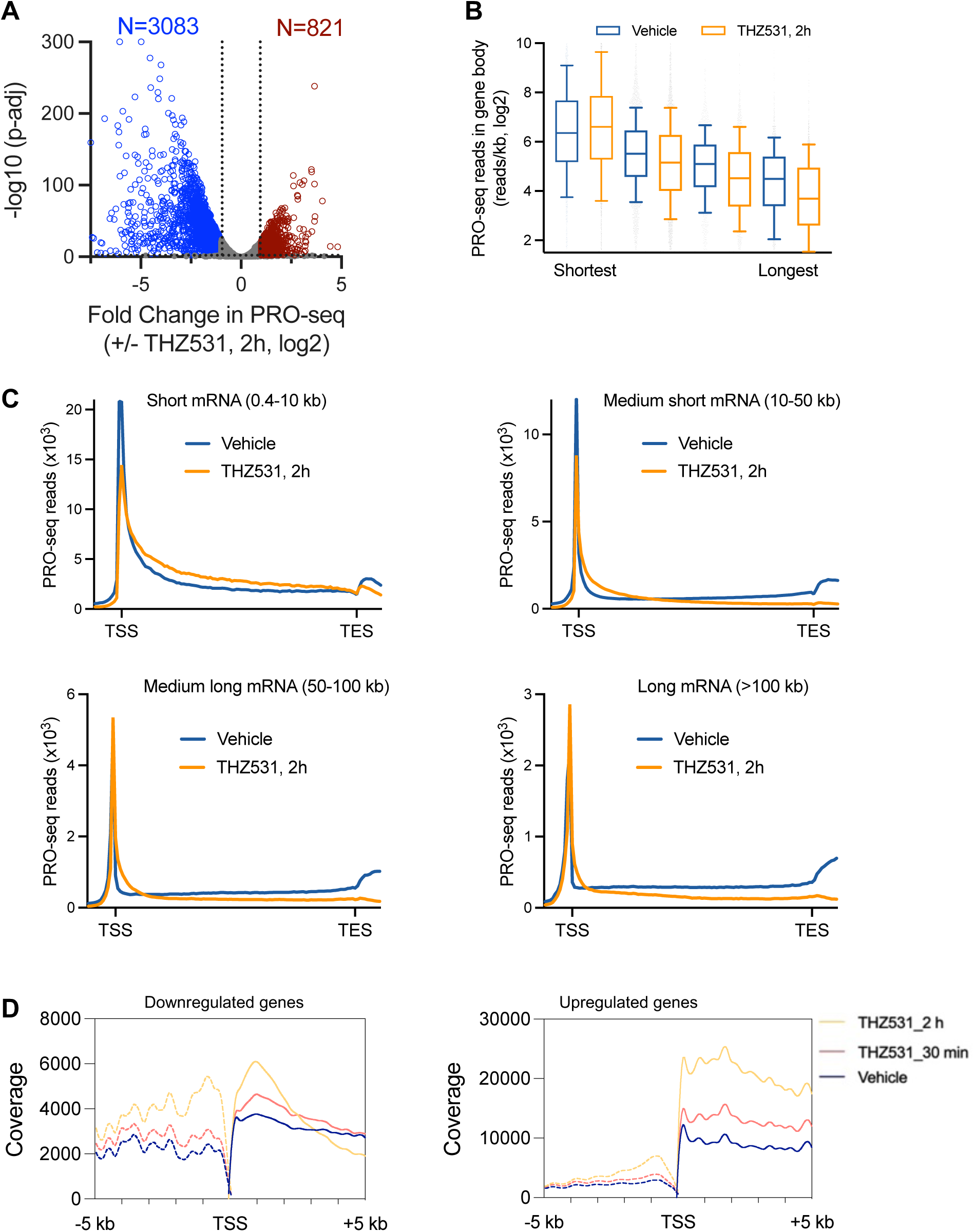
THZ531 treatment broadly disrupts Pol II elongation dynamics. A. Volcano plot shows fold changes and adjusted P-values (p-adj) from Precision Run-On sequencing (PRO-seq) data over the gene bodies of active mRNAs, comparing vehicle to THZ531 treatment. Affected genes (|log2FC| > 1; p-adj < 0.01) show both increased (red) and decreased (blue) Pol II signal upon THZ531 treatment, revealing differential effects on transcription. B. Boxplots depict PRO-seq read density in gene body region across all active mRNAs segregated into quartiles based on gene length. Boxes show 10th–90th percentiles and whiskers depict 1.5 times the interquartile range. C. Metagene plots of average PRO-seq signal in vehicle and THZ531-treated cells across gene groups segregated based on gene length. Bins from TSS to TES are scaled to gene length, with 100 bins/gene, and data outside gene bodies are shown as average reads per gene in 200-nt bins. D. Analysis of transient transcriptome sequencing (TT-seq) data from IMR-32 neuroblastoma cells treated with 400 nM THZ531 for 30 min, or 2 hours^15^. Coverage over the region 5000 bps up- or downstream of the TSS for genes that are 10-40 kb and up- (LFC > 1, q < 0.1) or downregulated (LFC < -1, q < 0.1) by THZ531 treatment. Increased reads within 1 kb of the TSS and also upstream of the TSS on the antisense strand indicate elevated transcription initiation for both groups of genes. However, downregulated genes exhibit a rapid decrease in reads following the 1 kb mark, which is indicative of decreased, productive elongation.

**Figure S8.**
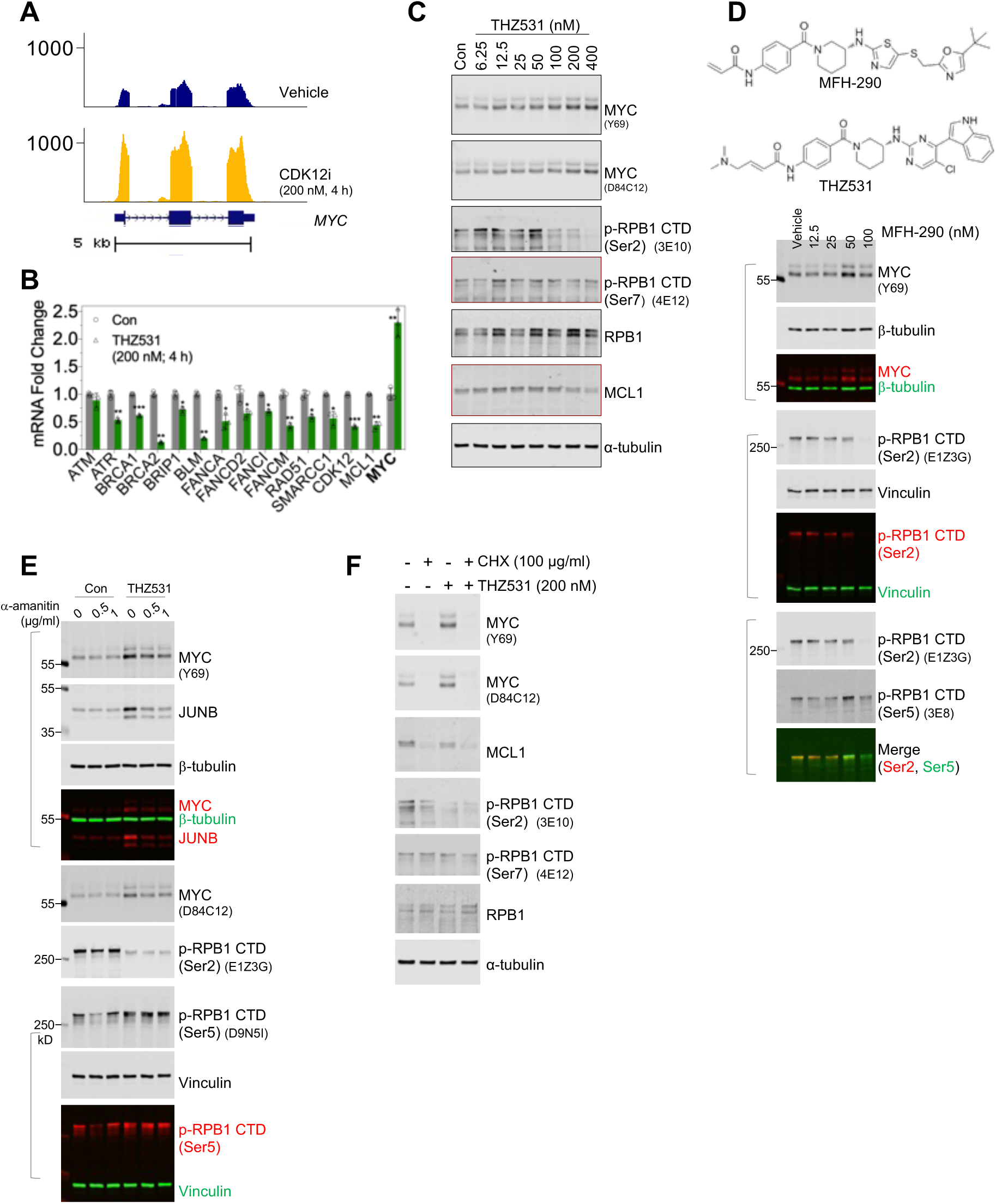
CDK12 inhibition upregulates MYC expression. A. Traces of RNA-seq reads for MYC in SKBR3 cells treated with vehicle control (0.4% DMSO, v/v) or THZ531 (200 nM) for 4 h. B. SKBR3 cells were as in (A) followed by total RNA extraction and reverse transcription. Quantitative PCR was performed for the indicated genes. Note that MYC expression is increased meanwhile other selected genes demonstrated significant downregulation. * p<0.05, ** p<0.01, and *** p<0.001 (Student’s t test). C. SKBR3 cells were treated with increasing doses of THZ531 for 4 hours. Cell lysates were prepared in SDS sample buffer and subjected to fluorescent immunoblotting. Clone numbers for monoclonal antibodies used are indicated. D. (top) chemical structures of MFH290 and THZ531. (bottom) HER2+ breast cancer cells HCC1954 were treated with vehicle or increasing doses of MHF-290 for 4 hours. Cells were then lysed with 1x SDS sample buffer, and cell lysates were subjected to fluorescent immunoblotting using the indicated antibodies. Note that 50 nM MFH-290 treatment reduced Ser2 CTD phosphorylation, but increased Ser5 CTD phosphorylation as well the protein abundance of MYC and JUNB. E. HCC1954 cells were treated as in (E) except the use of a-amanitin. The molecular weights of the fluorescent protein markers and clone identities for monoclonal antibodies are indicated. Merged images show signals from two primary antibodies raised in different species. F. THZ531-induced MYC expression relies on de novo protein synthesis. SKBR3 cells were treated with cycloheximide (100 μg/ml), or THZ531 (200 nM), either individually or in combination for three hours. Cell lysates were harvested for fluorescent immunoblotting.

**Figure S9.**
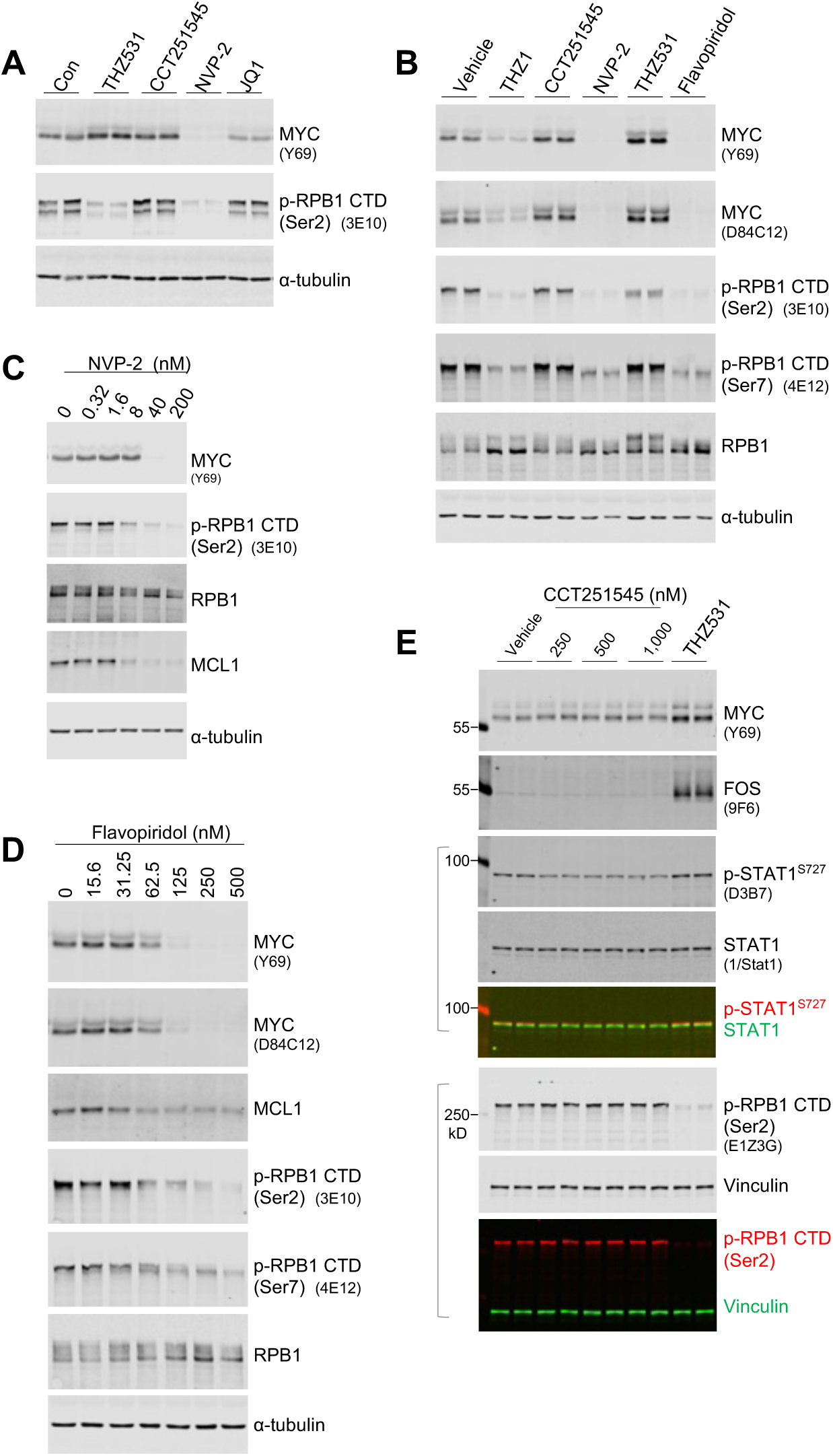
Effects of transcriptional CDKs inhibition on MYC expression. A. THZ531, but not inhibitors targeting CDK8 (CCT251545), or CDK9 (NVP-2), or BET family of bromodomain proteins (JQ1) increases MYC expression. SKBR3 cells were treated with indicated inhibitors (200 nM) for 4 hours, followed by fluorescent immunoblotting. B. HCC1954 cells were treated with indicated inhibitors (all 200 nM, except 500 nM for flavopiridol) for 4 hours, followed by fluorescent immunoblotting. C. HCC1954 cells were treated with NVP-2 for 4 hours at indicated concentrations. Note that NVP-2 does not obviously induce MYC expression at low doses, and totally abolished MYC expression at 40 or 200 nM. D. HCC1954 cells were treated with increasing concentrations of flavopiridol for 4 hours. Lysates were prepared and subjected to fluorescent immunoblotting. E. HCC1954 were treated with vehicle (0.08% DMSO, v/v), CDK8/19 inhibitor CCT251545 at the indicated concentrations, or THZ531 (400 nM). Four hours post treatment, cells were lysed with 1x SDS sample buffer, and cell lysates were subjected to fluorescent immunoblotting using the indicated antibodies. The molecular weights of the fluorescent protein markers and clone identities for monoclonal antibodies are indicated. Merged images show signals from two primary antibodies raised in different species. Note that for all immunoblotting based on short duration of treatment, cells were lysed with the same amount of sample buffer for complete lysis, and the same volume of lysates was loaded onto each lane of SDS-PAGE gels. Thus, each lane represents signal from similar number of cells.

**Figure S10.**
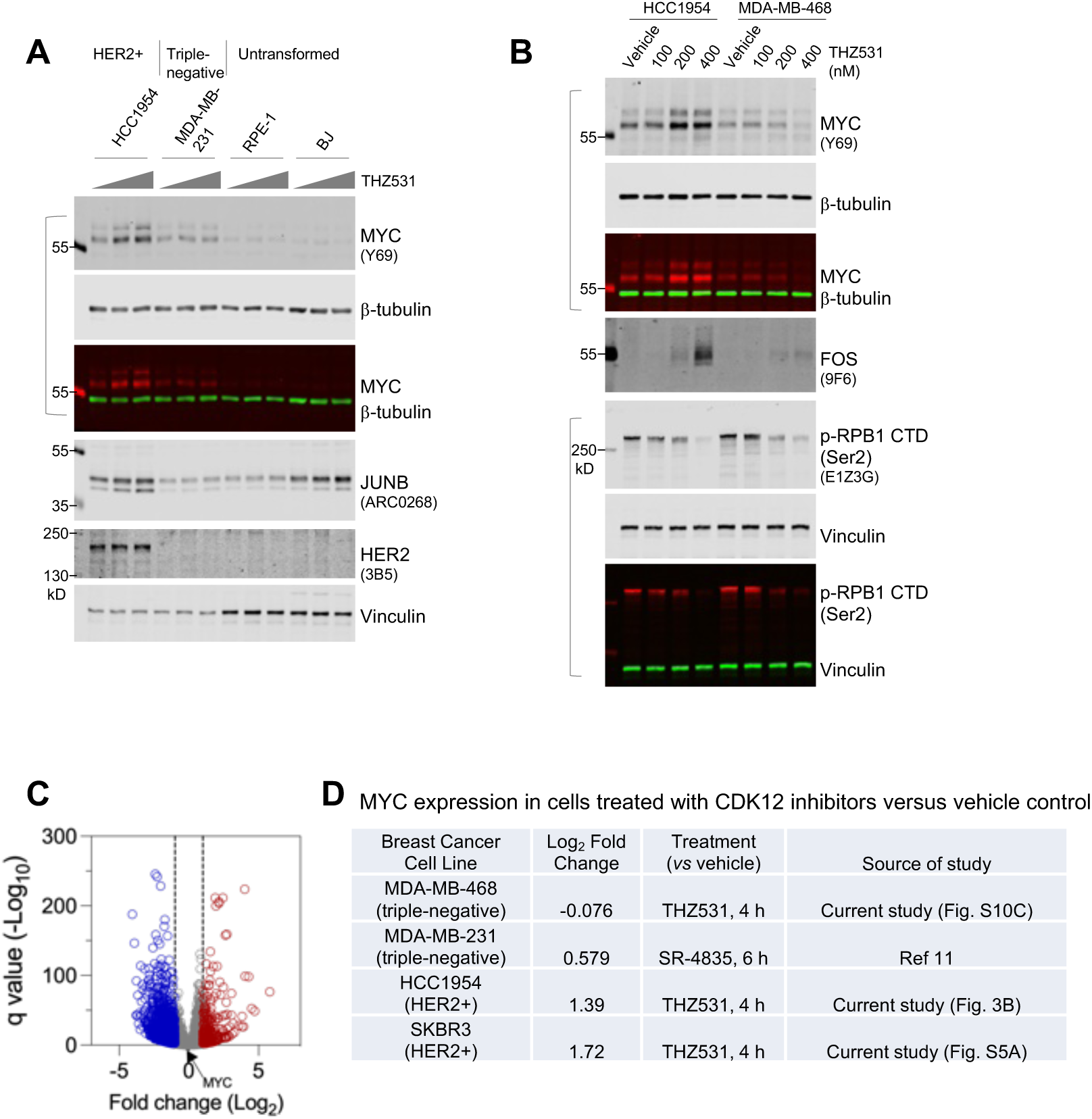
Effect of CDK12 inhibition on MYC expression. A. The indicated breast cancer cell lines and non-transformed human cells were treated with either DMSO vehicle control or THZ531 (200, 400 nM) for 4 h. Cells were then lysed using 1x SDS sample buffer and subjected to fluorescent immunoblotting. B. HER2+ breast cancer cells (HCC1954) and triple-negative breast cancer cells (MDA-MB-468) were treated with increasing doses of THZ531 for 4 h, followed by fluorescent immunoblotting. Note that FOS expression is induced in both cell lines, while MYC is only induced in HER2+ breast cancer cells. C. A volcano plot illustrates gene expression changes in triple-negative breast cancer cells (MDA-MB-468) treated with 200 nM THZ531 (4 h), compared with cells treated with vehicle control. Note that, despite the induction of a large number of genes, MYC expression is not altered by THZ531 treatment. D. A summary of MYC gene expression alteration following short-term exposure to CDK12 inhibitors. Note that robust MYC induction by CDK12 inhibition was also observed in the primary cultures of HER2+ metastatic breast tumor DFBC-1407 (Fig. 4C)

**Figure S11.**
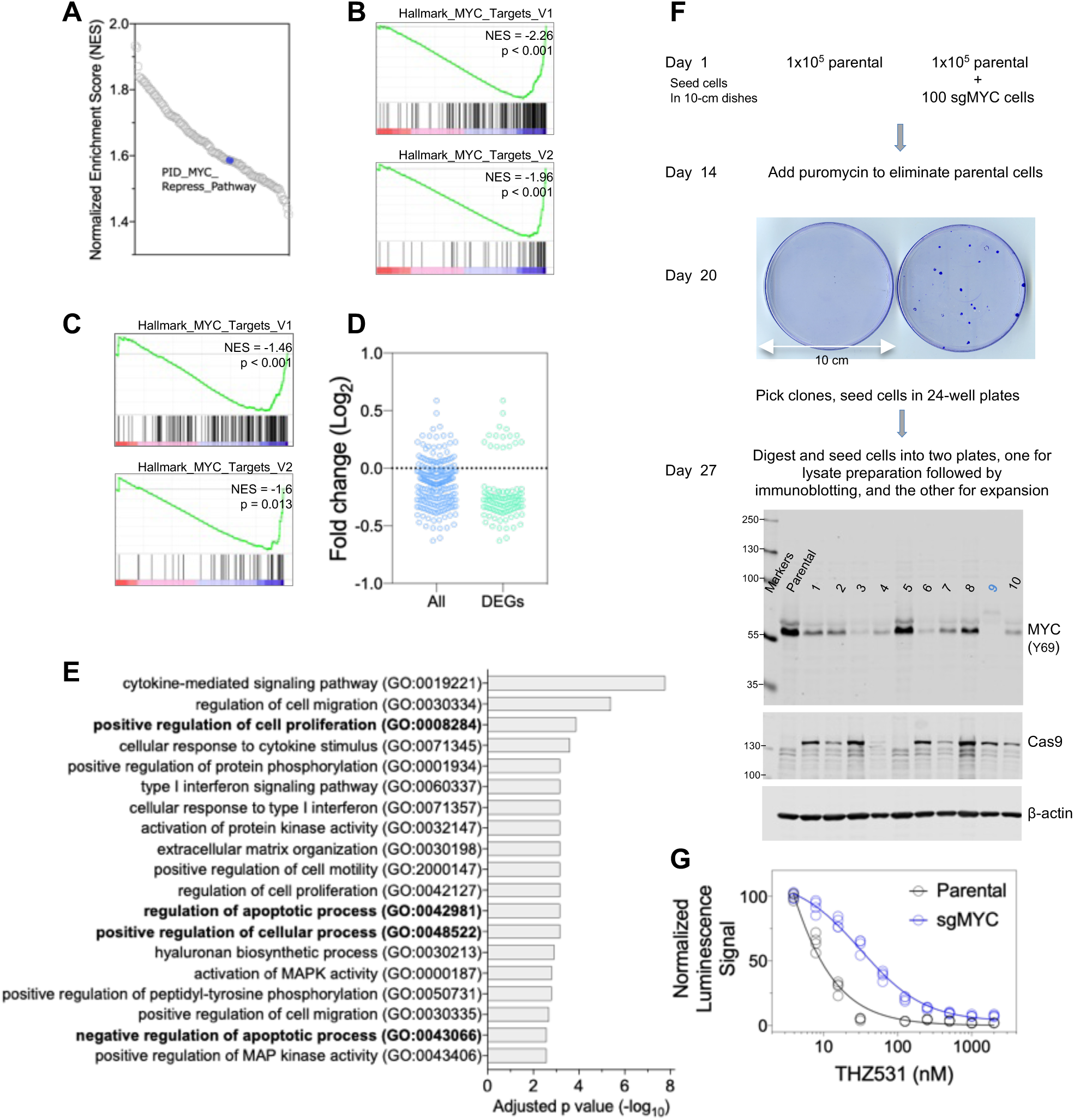
CRISPR/Cas9-mediated editing of MYC in HER2+ cancer cells alleviates the anti-proliferative potency of CDK12 inhibition. (A) The indicated MYC-mediated repression signature is upregulated in sgMYC cells treated with THZ531 compared to similarly treated parental controls. Summary plot of normalized enrichment scores (NES) for gene sets that are significantly upregulated in sgMYC HCC1954 cells treated with CDK12 inhibitor (nominal p-value < 0.05). (B) GSEA plots of indicated signatures in sgMYC HCC1954 cells treated with THZ531 (4 h, 200 nM) versus parental cells treated with THZ531. Normalized enrichment score (NES) and p values are indicated. (C) GSEA plots as in (B) for sgMYC HCC1954 cells in comparison to the parental controls. MYC signature is negatively enriched in sgMYC cells. (D) Expression analysis of the consolidated 235-gene MYC signature in sgMYC HCC1954 cells compared to parental controls. The first column shows the relative expression of all 235 genes, while the second column displays differentially regulated genes (DEGs, q < 0.1). (E) Overrepresentation analysis of GO_BP terms in the set of genes upregulated upon treatment with THZ531 in sgMYC HCC1954 cells compared to treated parental controls. Note that terms associated with the positive regulation of cell proliferation and the negative regulation of apoptosis are enriched. (F) Workflow outlining the construction of MYC knockout cancer cells. The HER2+ cancer line BT474 was transduced with sgMYC and subsequently seeded with parental cells at a low density. Puromycin was added in two weeks to eliminate parental cells. Outgrown clones were picked for expansion and validated by immunoblotting. Note that #9 clone in the blot was chosen as a knockout clone based on the absence of signals corresponding to the p64 and p67 forms of MYC (Hann et al.,1988). (G) MYC knockout clone (#9) and parental BT474 cells were treated with increasing concentrations of THZ531 in 96-well plates for 9 days before CellTiter-Glo assays.

**Figure S12.**
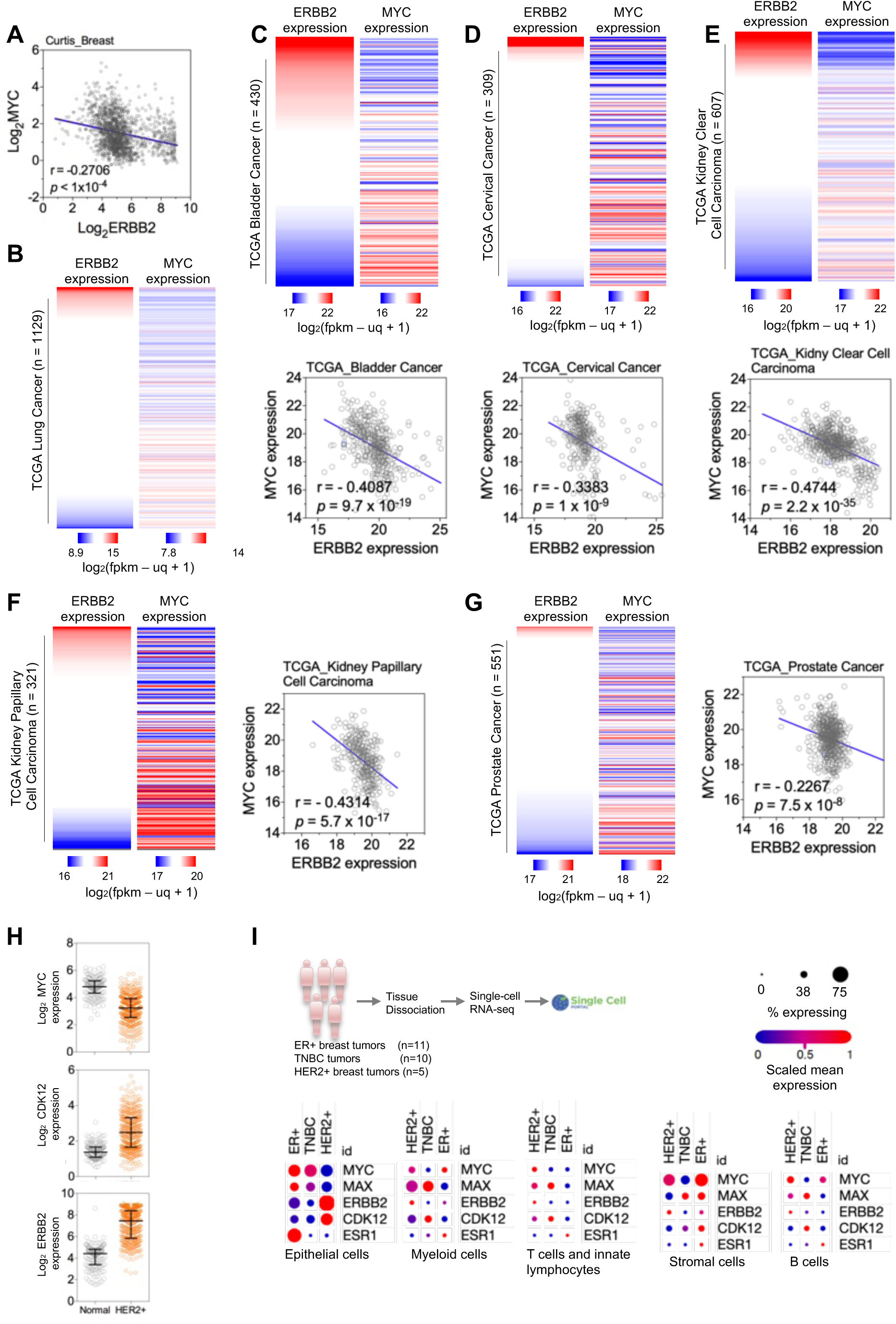
Tumors expressing high levels of ERBB2 display low MYC expression. A. MYC and ERBB2 gene expression data from the METABRIC cohort^60^ were obtained from Oncomine and analyzed using GraphPad Prism. The Pearson correlation coefficient (*r*) is indicated. B. Heatmap of ERBB2 and MYC expression in TCGA Lung Cancer dataset. The data were generated using the UCSC Xena Functional Genomics Browser (http://xena.ucsc.edu/). (C-G). Heatmaps and dot plots of ERBB2 and MYC expression in TCGA datasets for Bladder Cancer (C), Cervical Cancer (D), Kidney Clear Cell Carcinoma (E), Kidney Papillary Cell Carcinoma (F), and Prostate Cancer (G). RNA-seq data were obtained from the UCSC Xena Functional Genomics Browser and visualized using GraphPad Prism. H. Expression of MYC, CDK12, and ERBB2 in HER2+ breast cancer, compared to those in normal breast tissues. The gene expression data were generated by METABRIC cohort (*51*). Black lines in each group indicate median with interquartile range. I. (top) An overview of single-cell RNA-seq data generation and analysis. Single-cell RNA seq data of breast cancer from Wu et al. study^63^ were analyzed using the Broad Institute’s Single-Cell Portal (https://singlecell.broadinstitute.org/single_cell). (bottom) Heatmaps display the expression of indicated genes across various cell types in breast tumor samples. Note that epithelial cells within HER2+ breast cancer demonstrate the lowest expression of MYC and, as expected, the highest expression of ERBB2 and CDK12.

## Materials and Methods

### Antibodies

The following primary antibodies were purchased and used for fluorescence immunoblotting:

RNA polymerase II subunit B1 (phospho CTD Ser-2) Antibody, clone 3E10 (EMD Millipore, #04-1571);

Phospho-Rpb1 CTD (Ser2) (E1Z3G) Rabbit mAb (Cell Signaling Technology, #13499);

Phospho RNA Polymerase II (S2) Antibody, (Bethyl Laboratories, A300-654A);

RNA polymerase II subunit B1 (phospho-CTD Ser-5) Antibody, clone 3E8 (EMD Millipore, #04-1572);

Phospho-Rpb1 CTD (Ser5) (D9N5I) Rabbit mAb (Cell Signaling Technology, #13523)

RNA polymerase II subunit B1 (phospho-CTD Ser-7) Antibody, clone 4E12 (EMD Millipore, #04-1570);

RNA Polymerase II Antibody (Bethyl Laboratories, A300-653A);

RNA Polymerase II RPB1, clone 8WG16 (BioLegend, #664906);

Rpb1 NTD (D8L4Y) Rabbit mAb (Cell Signaling Technology, #14958);

c-Myc (clone Y69) Rabbit mAb (Abcam, #ab32072);

c-Myc (D84C12) Rabbit mAb (Cell Signaling Technology, #5605);

anti-CDK12 (Cell Signaling Technology, #11973);

anti-CDK12 (proteintech, #26816-1-AP);

anti-CDK12, clone 45F7-H2 (BIO-RAD, #VMA00874);

anti-CDK13, clone 46B7-G7 (BIO-RAD, #VMA00875);

CDK7 Recombinant Monoclonal Antibody (BL-80-5D4) (Bethyl, #A700-006);

CDK7 (MO1) Mouse mAb (Cell Signaling Technology, #2916);

CDK8 (D6M3J) Rabbit mAb (Cell Signaling Technology, #17395);

CDK9 (C12F7) Rabbit mAb (Cell Signaling Technology, #2316);

Anti-CDK19 (Sigma-Aldrich, #HPA007053);

JunB Rabbit mAb, clone ARC0268 (ABclonal, #A4848);

c-Jun (60A8) Rabbit mAb (Cell Signaling Technology, #9165);

c-Fos (9F6) Rabbit mAb (Cell Signaling Technology, #2250);

Anti-c-ErbB2/c-Neu (Ab-3) Mouse mAb (3B5) (EMD Millipore, #OP15);

anti-PARP (Cell Signaling Technology, #9542);

anti-β-Actin, clone AC-15 (Sigma, A5441);

anti-Vinculin (Sigma, V9131); anti-β-Tubulin (BioLegend, # 903401).

Secondary antibodies include the following: Alexa Fluor 680 Goat Anti-Rat IgG (Life Technologies, #A21096), Alexa Fluor 680 Goat Anti-Rabbit IgG (Life Technologies #A27042), and DyLight 800 conjugated anti-mouse IgG (RockLand, #610-445-002).

### Chemicals

THZ531, MFH-290, THZ1, and NVP-2 were the generous gifts of Nathanael Gray’s group. Commercially purchased chemicals are as following: α-amanitin (MedChemExpress, #HY-19610); actinomycin D (Cell Signaling Technology, #15021); doxorubicin (Sigma, #D1515); pan Caspase Inhibitor Z-VAD-FMK (R&D, #FMK001); SR-4835 (TargetMol, #T8325).

### Fluorescent Immunoblotting

Upon harvest, cells were rinsed with PBS, and lysed immediately with 1x SDS sample buffer consisting of 50 mM Tris–HCl, (pH 6.8), 2% (w/v) sodium dodecyl sulfate (SDS), 0.04% (w/v) bromophenol blue, 10% (v/v) glycerol, and 5% (v/v) β-mercaptoethanol. The typical amount of sample buffer used in one well of a 12-well plate seeded with 0.2 million cells the prior day is 200 μl. Cell lysates were collected and boiled to achieve complete denaturation of protein and sheering of genomic DNA. Typically, 20 μl lysates were loaded into each lane of a freshly casted sodium dodecyl sulphate–polyacrylamide gel electrophoresis (SDS-PAGE) gel, and electrophoresis was performed with ice-water incubation for heat dissemination. Proteins were then transferred onto nitrocellulose membrane (Thermo Scientific, #88018). The membrane was blocked with Blotting Grade Blocker Non Fat Dry Milk (5%, w/v; Bio-Rad, #1706404XTU), and then incubated with primary antibodies diluted 1:1000 fold. Following overnight incubation at 4°C, the membrane was washed, and incubated with fluorescent-dye conjugated secondary antibodies (1:5000 fold dilution) for 1 h at room temperature. Following extensive washes, the membranes were scanned using an Odyssey CLx Near-Infrared Fluorescence Imaging System according to the manufacturer’s instructions. The images were then exported as TIFF files and processed in ImageJ (NIH) only for the basic operations of cropping and rotation.

For all immunoblotting based on short-term treatment experiments (*e.g.,* 4 h), the same volume of lysis buffer was added to each treatment group, and the same volume of lysate was loaded into each lane of SDS-PAGE gels. Given that short-term treatment of cancer cells with inhibitors does not alter the total cell number, the signal of fluorescent immunoblotting can be regarded as that *per* cell.

### Immunofluorescence

Cells were harvested via trypsinization and then seeded on No. 1.5 coverslips (12 mm diameter) pre-placed into 12-well plates at a density of 2x10^5^ cells *per* well. Following overnight incubation, cells were refreshed with medium containing vehicle control (DMSO, 0.8%, v/v), THZ531 (400 nM), NVP-2 (200 nM), or flavopiridol (400 nM). Four hours post treatment, cells were fixed with 4% formaldehyde for 10 min. After washing, cells were permeabilized with 0.1% Trition X-100 for 10 min. Cells were then washed and blocked with 1% bovine serum for 30 min before incubation with the primary antibodies prepared in PBS containing 1% bovine serum albumin. After overnight incubation at 4°C, the samples were washed and incubated with fluorophore-conjugated secondary antibodies for 1 h at room temperature. After extensive washing, the samples were mounted with ProLong Diamond Antifade Mount (Life Technologies, # P36965). Imaging acquisition was performed using NIS-Elements image acquisition software and with Yokogawa W1 spinning disk confocal on an inverted Nikon Ti fluorescence microscope (Nikon Imaging Center, Harvard Medical School). Fiji was used for image analysis, including merging channels with different colors and cropping. The primary antibodies used include the following: Phospho-Rpb1 CTD (Ser2) (E1Z3G) Rabbit mAb (Cell Signaling Technology, #13499); Phospho-Rpb1 CTD (Ser5) (D9N5I) Rabbit mAb (Cell Signaling Technology, #13523); mouse monoclonal Anti-EMD/Emerin antibody, clone CL0203 (Sigma-Aldrich, #AMAB90562). Fluorophore-conjugated secondary antibodies include Alexa Fluor 488 anti-rabbit IgG (H+L) (Life Technologies, #A27034), and Alexa Fluor 647 anti-mouse IgG (BioLegend, #405322).

### CRISPR/Cas9-mediating gene editing Construction of all-in-one lentiCRISPR vectors

All-in-one lentiCRISPR v2 vector (Sanjana et al., 2014) was obtained via Addgene (#52961) and used to introduce guide sequences targeting GFP (as a control), MYC, or transcriptional CDKs. Forward and reverse oligos were synthesized at Eton Bioscience and used for annealing, followed by ligation with a BsmBI-digested lentiCRISPR backbone. The ligation reaction was then transformed into competent E. coli (stbl3 strain) and followed by overnight incubation in a warm room. Single clones were picked for PCR-based identification using a U6 promoter paired with the reverse oligo of the guide inserts.

Positive colonies were then further cultured for MidiPrep (Qiagen, #12943). Plasmids were verified by sequencing using the U6 primer available at the Eton Bioscience facility. Sequences of oligos are as following (note that the part following CACCG in the forward sequence is gene-specific):

sgGFP_fwd: CACCG GGGCGAGGAGCTGTTCACCG

sgGFP_rev: AAAC CGGTGAACAGCTCCTCGCCC C

sgMYC_fwd: CACCG GCCGTATTTCTACTGCGACG

sgMYC_rev: AAAC CGTCGCAGTAGAAATACGGC C

sgCDK12_1_fwd: CACCG GGCAAGAAGGACGGGAGTGG

sgCDK 12_1_rev: AAAC CCACTCCCGTCCTTCTTGCC C

sgCDK 12_2_fwd: CACCG AGGGAGAACGACGAACGTCG

sgCDK 12_2_rev: AAAC CGACGTTCGTCGTTCTCCCT C

sgCDK7_fwd: CACCG GAAGCTGGACTTCCTTGGGG

sgCDK7_rev: AAAC CCCCAAGGAAGTCCAGCTTC C

sgCDK8_fwd: CACCG CGAGGACCTGTTTGAATACG

sgCDK8_revrv: AAAC CGTATTCAAACAGGTCCTCG C

sgCDK8_fwd: CACCG GCACCGCAAGACCGGCCAGA

sgCDK8_rev: AAAC TCTGGCCGGTCTTGCGGTGC C

sgCDK9_fwd: CACCG GCACCGCAAGACCGGCCAGA

sgCDK9_rev: AAAC TCTGGCCGGTCTTGCGGTGC C

sgCDK13_fwd: CACCG AGGAGCGGCAACAGCAGCGG

sgCDK13_rev: AAAC CCGCTGCTGTTGCCGCTCCT C

sgCDK19_fwd: CACCG ATTATGCAGAGCATGACTTG

sgCDK19_rev: AAAC CAAGTCATGCTCTGCATAAT C

### Packaging virus for transducing cancer cells

HEK293T cells were used for lentivirus packaging. Typically, cells were seeded in T-25 tissue culture flasks at the density of 2.5 - 3 million cells per flask. On the second day, 4 µg DNA (2 µg vector DNA, 1.5 µg pCMVdR8.91, and 0.5 µg pMD2-VSVG) and 12 µl polyethylenimine (PEI; homemade with powder purchased from Polysciences, # 23966-2) were diluted in 300 µl PBS (Gibco, #10010023), mixed, and then incubated at room temperature for 15 min. The lipid/DNA mixture was added to HEK293T cells. In 24 hours, transfected cells were refreshed with new medium, while the cells to be transduced were seeded in 6-well plates. Forty-eight and then seventy-two hours post the initial transfection, viral supernatant was collected and filtered through 0.45-µm filters and immediately used for transducing target cells supplemented with polybrene at a final dose of 8 µg/ml (Millipore, # TR- 1003-G). Forty-eight hours after the initial infection, transduced cells were subjected to 1-2 days of puromycin selection, followed by assays to examine the efficiency of gene editing and potential effects on cell growth and/or response to drug treatment.

### Clonal expansion of sgMYC cells

Following transduction and puromycin selection, BT474_sgMYC cells were seeded at the density of 100 cells per 10-cm dish together with 0.1 million parental BT474 cells. Upon reaching visual confluency, cells were refreshed with medium containing puromycin (1 µg/ml) to eliminate non-transduced cells. The cells were cultured until individual clones had reached a size sufficient for manual picking under a 4x objective. Clones were picked into 24-well plates and, in 1-2 weeks, digested for immunoblotting and, therefore, examination of MYC protein.

### Cell growth assays

Cells were trypsinized and resuspended in medium to measure cell density using the Countess Automated Cell Counter (Life Technologies). Cell suspensions of various densities, ranging from 5,000 to 20,000 cells per ml, were prepared and seeded in multiple-well plates. For clonogenic assays, each well was replenished with medium 4-5 days after seeding. At assay endpoints, cells were fixed with formalin and subsequently stained with crystal violet. The stained plates were scanned before the staining was extracted using 10% acetic acid and absorbance measured at a 570 nm wavelength (750 nm used as the reference).

For assays performed in 96-well plates, 100 µl cell suspension was seeded into each well, followed by the addition of an equal volume of medium consisting of increasing concentrations of drug. Upon harvest, CellTiter-Glo 2.0 Cell Viability Assay (Promega, #G9241) was used to determine the ATP level in culture, based on the instructions provided by the manufacturer. Briefly, following addition of CellTiter-Glo mixture into wells, cell lysates were incubated at room temperature for 15 min with shaking, and then transferred into black-walled plates for reading of luminescence values (GloMax Navigator Microplate Luminometer, Promega).

### Cell culture

HER2+ cancer cells were purchased from the American Type Culture Collection (ATCC), and were cultured with RPMI1640 (Invitrogen, #11875) supplemented with 10% (v/v) fetal bovine serum and Penicillin-Streptomycin (Gibco, #15140122; final dose is 100 units *per* ml). Cells were incubated at 37 °C with 5% CO2 and passaged twice a week using Trypsin-EDTA (0.05%) (Invitrogen, #25300). Cells were tested for mycoplasma contamination using the MycoAlert Mycoplasma Detection Kit (Lonza, #LT07-318), according to the manufacturer’s instructions.

### Primary culture of HER2+ metastatic breast cancer

DFBM-NI17 primary culture was previously established (Ni et al., 2022), and was maintained in NeuroCult NS-A Proliferation Kit (Human) (STEMCELL Technologies, #05751) with the supplementation of heparin (2 μg/ml), EGF(20 ng/ml), and bFGF (10 ng/ml). DFBM-NI17 was maintained in suspension using ultra-low adherence tissue culture vessels.

To derive primary culture of DFBC-1407, passage 1 patient-derived xenograft (PDX) tumors of DFBC- 1407 (Goel et al., 2016) were minced and resuspended in Collagenase/Hyaluronidase (STEMCELL Technologies, #07912), followed by incubation at 37°C with rotation for 2 h. The cells were then recovered by centrifugation and seeded in the DMEM/F-12 medium supplemented with 5% horse serum (Life Technologies, #16050122), 1% penicillin/streptomycin, 20 ng/ml EGF, 0.5 mg/ml hydrocortisone, 100 ng/ml cholera toxin, and 10 μg/ml insulin. When cells grew and reached confluence, they were harvested for storage in liquid nitrogen as well as passaging and experiments. Unlike DFBM-NI17, the primary culture of DFBC-1407 attaches to vessel surface and present epithelial cell morphology.

### Xenograft tumor models and drug efficacy studies

All animal studies were conducted in accordance with the animal use guidelines from the National Institutes of Health, and animal protocols were approved by the Dana-Farber Cancer Institute Animal Care and Use Committee. HER2+ breast cancer cells (HCC1954) were harvested and resuspended in 40% Matrigel-Basement Membrane Matrix (Corning, #CB-40234) to reach a density of 10 million cell *per* ml. The cell suspension was injected into the third pair of mammary fat pads of SCID mice (100 μl *per* site). When tumors reached approximately 200 mm^3^ in size, mice were randomly divided into two groups: one for vehicle control and other for the administration of the CDK12 inhibitor SR-4835. For the preparation of vehicle control, DMSO was mixed at a ratio of 1:9 with 30% HP-β-CD (2-Hydroxypropyl-β-cyclodextrin; MedChemExpress, #HY-101103) dissolved in water. The CDK12 inhibitor SR-4835 (TargetMol, #T8325) was dissolved in DMSO and aliquoted; upon use, the SR-4835 solution was mixed at a ratio of 1:9 with 30% HP-β-CD. Vehicle and SR-4835 were administered by oral gavage 6 days *per* week at 20 mg/kg. Tumors were measured twice a week using a caliper, and tumor volume was calculated using the formula: tumor volume = 0.5 × length × width × width.

### Fluorescence recovery after photobleaching (FRAP)

FRAP were performed as previously described (Steurer et al., 2018). For FRAP analysis in GFP-RPB1 knockin cells, a Leica SP5 confocal microscope using a HCX PL APO CS 63x, 1.40NA oil-immersion lens and LAS AF software. Cells were placed in a controlled environment with 37°C and 5% CO2 during the experiments. Fluorescence of GFP-RPB1 was detected using a 488 nm argon laser. At pixel size 24.6 × 24.6 μm, a strip of 512 × 32 pixels spanning the nucleus was imaged every 400 ms at 400 Hz. Twenty-five frames were recorded before the bleach pulse. GFP fluorescence in the strip was bleached for one frame with 90% laser power. The recovery of fluorescence was monitored for 200 seconds (500 frames) within and outside the strip. FRAP curves were corrected for background fluorescence outside the nucleus and normalized to prebleach fluorescence in the region of interest, which was set at 1. Cells were either untreated or treated with 400 nM THZ531 for 90 min, or 2 µM THZ1 for 90 min prior FRAP analysis.

### Precision Run-on sequencing (PRO-seq)

For PRO-seq, 5 million HCC1954 cells were seeded in 15-cm dishes 18 hours prior to treatment. The cells were treated with DMSO or THZ531 (N=2, 2 h) and permeabilized as described (Mimoso and Goldman, 2023). To the aliquots of 1 million permeabilized HCC1954 cells, 100,000 permeabilized *Drosophila* S2 cells were added prior to the run-on reaction based on cell count. Run-on reaction and library prep from the resulting RNA was performed as described (same citation as above) and the subsequent libraries were sequenced on Novaseq 6000 using a high output flow cell for 100 cycles in paired ended fashion.

FASTQ files were trimmed to 100 bp and read pairs with average base quality scores below 20 were removed. Adaptor sequences along with low-quality reads were then removed using cutadapt (Martin 2011), and any subsequent reads shorter than 20 nt were discarded. The last 3’ nucleotide was deleted from each trimmed read followed by mapping using bowtie (1.2.2) to the *Drosophila dm6* genome (-k1 - v2 -best -X100 –un) to measure the spike return per samples. Next, the resulting unaligned reads were mapped to the hg38 reference genome using the same parameters. Uniquely aligned read pairs were extracted, and single-nucleotide resolution bedGraph files were generated based on 3’ end mapping positions. After comparing the reads aligned to the *hg38* and *dm6* genomes, we observed a consistently lower percentage of sequenced reads mapping to the spike-in *dm6* genome in THZ531 treated samples across the replicates, indicative of a modest decrease in transcription in THZ531 treated cells. Spike normalization ratios were thus calculated for each sample by dividing the number of reads mapped to *dm6* genome by the numbers of reads corresponding to the sample with the fewest *dm6* mapped reads (1.13 and 1.0 for replicates belonging to control; 1.33 and 1.26 for replicates belonging to THZ531-treated condition, respectively). The final reads were then scaled by dividing the number of reads mapped to *hg38* by the spike ratio using normalize_bedGraph (Martin et al., 2021) and merged using bowtie2stdBedGraph.pl (Martin et al., 2021), followed by generation of binned bedGraphs in 25 bp windows.

### Gene Annotation and Transcriptional Unit Definition

Gene annotation was performed using the Get Gene Annotation (GGA) pipeline (https://github.com/AdelmanLab/GetGeneAnnotation_GGA) to define dominant transcriptional start sites (TSS) and transcriptional end sites (TES) for each gene. The GGA analysis integrated PRO-seq data with reference genome annotations (hg38) to identify dominant TSS positions based on PRO-seq signal intensity, transcriptional end sites defined by 3’ PRO-seq signal termination, and gene boundaries encompassing the full transcriptional unit from dominant TSS to TES. Genes shorter than 400 base pairs were excluded from downstream analyses to ensure robust signal detection. The final gene annotation file contained 13,531 transcriptional units, with 11,876 protein-coding genes meeting length criteria for detailed analysis. PRO-seq-based differential expression was performed using DESeq2 on read counts spanning from TSS+250bp to TES for each gene. This approach focused on gene body transcription while excluding promoter-proximal signals. Spike-in normalization factors were incorporated as custom size factors in DESeq2. Sample-specific normalization factors included Veh_2h_R1 (1.13), Veh_2h_R2 (1.00), THZ531_2h_R1 (1.33), THZ531_2h_R2 (1.26). Significance thresholds were set at |log2FC| > 1.0 and adjusted p-value < 0.01. The PRO-seq data generated in this study has been deposited in the Gene Expression Omnibus (GEO) database (accession no. GSE302011). The following secure token has been created to allow review of record GSE302011 while it remains in private status: kjopqwyonjovxut.

### Enhancer Classification and Annotation

Enhancer identification was performed using dREG (discriminative Regulatory Element detection from GRO-seq) analysis (Danko et al., 2015) on combined PRO-seq data. For dREG, forward and reverse 3’ PRO-seq bedGraph files were merged across all samples and converted to bigWig format for dREG input. dREG was run using the pre-trained support vector machine model (asvm.gdm.6.6M.20170828.rdata) with default parameters (Filtering criteria: dREG score ≥ 0.4; adjusted p-value ≤ 0.026). dREG peaks within 1kb of dominant TSSs were removed to focus on enhancer elements, resulting in 179,120 promoter-distal peaks from an initial 202,423 filtered peaks. Peaks overlapping problematic repetitive elements (rRNA, tRNA, snRNA, scRNA, srpRNA) were removed using UCSC RepeatMasker annotations, eliminating 808 additional peaks. Remaining peaks (178,304) were classified as Intergenic enhancers (137,274 peaks located outside annotated gene bodies) or Intragenic enhancers (40,956 peaks within gene bodies but on the opposite strand relative to the host gene). Next, dREG peaks were intersected with TSS call output to identify enhancers with detectable transcriptional initiation (Henriques et al., 2018). Strand orientation was assigned for TSS-overlapping enhancers based on the dominant TSS strand from TSS call analysis; dREG peaks not overlapping with TSScall output were discarded.

### Super Enhancer Identification

Position based clustering analysis identified super enhancers using TSS clustering analysis as defined earlier (Whyte et al., 2013). High-activity thresholds classified genes associated with >30 clustered TSSs as super enhancer-associated. Functional validation of super enhancer-associated genes was performed through transcriptional activity analysis using PRO-seq signal analysis across gene bodies (TSS+250bp to TSS+5kb). Chromatin state analysis utilized H3K27Ac ChIP-seq data from HCC1954 cells processed (GEO: GSE72956) using a standard ChIP-seq pipeline. The pipeline included alignment to hg38 genome using bowtie, duplicate removal and quality filtering, 25bp binning for signal quantification, and peak calling with signal normalization. Enhancer-chromatin correlation involved profiling H3K27Ac signal across all identified enhancer regions (Martin et al., 2021) with 4kb flanking regions (±4kb from enhancer center) to assess chromatin accessibility and active enhancer marks. Comparative analysis employed Wilcoxon rank-sum tests to assess statistical significance for gene length distributions across expression categories, PRO-seq signal intensities at enhancers and super enhancer-associated genes, and treatment-induced changes in enhancer activity.

### RNA Extraction and Sequencing

Cells were treated for 4 hours with the chemicals indicated in the figure legends, and total RNA was purified using the Monarch Total RNA Miniprep Kit (New England Biolabs, #T2010). On-column DNase digestion was performed to remove any residual DNA. The integrity of total RNA was assessed by an Agilent Bioanalyzer 2100 at Novogene. The cDNA library was constructed using NEBNext® Ultra™ II RNA Library Prep with Sample Purification Beads (New England Biolabs, #E7775). Briefly, a total of 400 ng RNA per sample was subjected to poly(A)-selection, followed by reverse transcription to generate cDNA. The resulting cDNA was subjected to adapter ligation and PCR amplification. Library quality control was performed using Labchip and qPCR. Sequencing was then performed on a Novaseq 6000 to generate 150 bp paired-end reads (Novogene).

### RNA Sequencing Analysis

Raw reads containing adapters, base quality scores less than 5 and more than ten percent N’s were removed using Cutadapt v1.16. Processed reads were then mapped to hg19 with STAR v2.7.3a 2-pass alignment, first using a genome index with 100 bp overhangs and then using an index built with 75 bp overhangs and a combination of the junctions determined from the initial alignments of control and treated duplicates. Unstranded gene hits were then counted for both alignments using HTSeq v0.9.1 with a minimum alignment quality of 10 and differential expression was determined in DESeq2 using a beta prior. All genes having an average TMM-normalized CPM less than 1 were removed. Gene IDs were then annotated to their gene symbols and lowly expressed duplicates were also removed based on their average CPM. As independent validation, gene-level differential expression was determined from transcript abundances quantified in Kallisto v0.46 using the hg19 transcriptome. The differential expression analysis results of STAR and Kallisto were evaluated for correlation.

The RNA-seq data generated in this study has been deposited in the Gene Expression Omnibus (GEO) database (accession no. GSE181905).

For published datasets (PRJNA530774, GSE113313, GSE104837, GSE116282, GSE116017, GSE108615, GSE80409) NCBI SRA-normalized reads were downloaded using fastq-dump and analyzed with Kallisto v0.46. Hg19 was used as the transcriptome assembly. Transcript abundances were quantified by applying default parameters and the strandedness information in the associated publication. For single-end reads, an average fragment length of 200 and standard deviation of 20 was used, unless otherwise specified. Gene-level differential expression was then determined in DESeq2 using a logarithmic fold change prior (Love et al., 2014).

All volcano plots were plotted in PRISM using the log2 fold-change and adjusted p values determined with DESeq2. Significantly altered genes are those with |log2 FC| > 1 and q < 0.1. Genes were also annotated to their longest transcript and the size distributions of significantly down or upregulated genes were compared using a Wilcoxon signed-rank test. For gene set enrichment analysis, lists containing all expressed genes ranked by both the direction and statistical significance of their differential expression were fed to GSEA v4.1.0 using default parameters and the Hallmark, Kegg, PID, Reactome and Biocarta pathway or gene set collections. Several other gene sets containing MYC targets and DNA damage response genes were also assessed for statistically significant enrichment.

### Section for RNA-seq trace

The pre-processed reads were aligned to hg19 now using ENCODE parameters and quantitated as above using Kallisto and HTSeq. The new alignments were confirmed to be correlated with the previous alignment data. The biological duplicates of each individual treatment condition were then evaluated for correlation. This was done first using the Kallisto TPM values of all expressed transcripts for the filtered set of genes and then using the TPM values derived from the gene-level HTSeq counts. Since the correlation was high, the alignment files of the two replicates were coordinate-sorted and then merged using SAMtools v1.9. The bamCoverage function in deepTools v3.5.0 was then run to generate bigwig files of the merged alignments by dividing the genome into bins of 20 bps and normalizing coverage to 1x. The bin counts over specifIc gene loci were determined for the individual treatment conditions using the multiBigWigSummary function in deepTools and then plotted in Prism.

### Metagene profiling

Expressed genes with a |LFC| > 0.5 and q < 0.1 were determined for each of the treatment conditions (K12i, K9i, K7i) and then intersected. The intersected set of genes were then segmented based on size (2.5 – 10 kb; 10 – 50 kb; 50 – 100 kb; > 100 kb). These were then used to subset the hg19 transcriptome and generate new gtf files, which were used as metagene models for the computeMatrix function in deepTools. computeMatrix was run on the previously generated bigwig files by scaling the gene body to 500 bp regions with an unscaled 100 bp region up- and downstream and then averaging the score over the regions in 5 bp bins. The mean profile of the score distributions for each differently-sized gene set was determined using the plotProfile function in deepTools and then plotted in Prism.

### TCGA Data Analysis

Raw counts were obtained from the Broad GDAC Firehose and UCSC Xena Data Browser for the Ovarian, Prostate, and Breast Cancer cohorts. Counts were TMM-normalized and log-transformed using edgeR v3.13. PAM50 genes were selected and the resulting matrices were used to generate tSNE plots using Rtsne v0.15 with a perplexity of 40. To define CDK12 high or low groups, samples having normalized CDK12 expression that is in the top or bottom 5% of all samples were determined. The raw counts of these two groups were normalized and then manipulated to determine differential enrichment using DESeq2 with a beta prior. The same analysis was performed based on CDK7 and CDK9 expression. Lowly expressed genes and duplicates were removed as before and volcano plots were generated in PRISM.

### Consolidating a MYC signature

To define a MYC signature, 12 different gene sets associated with MYC activity, including the Hallmark MYC_v1, and v2 sets, were merged. The top correlates of MYC gene expression were determined using the TPM values of 19144 genes in 1305 cancer cell lines available from DepMap (20Q2). These were defined as the positive correlates located above the point with the greatest change in slope following the inflection point. The 1246 highly correlated genes were interested with those from the merged gene sets to derive a MYC signature having 235 genes, of which 77 are shared with the two hallmark sets. The differential enrichment of this signature was evaluated for each CDK low vs high comparison. Gene list of the consolidated MYC signature is included in the supplemental file.

### Alternative Polyadenylation

As an alternative to 3’-seq, was used to determine alternative polyadenylation. QAPA is similar to 3’-Seq in that it relies on a database of 3’ UTRs containing polyadenylation sites refined by integrating numerous 3’-Seq datasets. In this case, the human genome hg19 was used to assemble the database of 3’ UTRs with distal and proximal polyadenylation sites. Their locations were determined and appended to an hg19 transcriptome gtf file. Salmon v1.3.0 was then used to quantify the different 3’ UTR isoforms. Specifically, an index was built with Salmon using a kmer length of 31 and a dense hash. Transcript abundances were then determined from the processed reads according to the assembled index. With QAPA, genome-wide distal and proximal polyadenylation usage was determined from the expression of the different 3’ UTR isoforms in each sample. Based on this information, the change in proximal polyadenylation usage (ΔPPAU) was determined genome-wide.

The Welch’s t-test was then used to determine the statistical significance of the difference between PPAU in the treated and control duplicates and define genes with either proximal or distal polyadenylation, or no change.

